# Vascular Progenitors Generated from Tankyrase Inhibitor-Regulated Naïve Diabetic Human iPSC Potentiate Efficient Revascularization of Ischemic Retina

**DOI:** 10.1101/678599

**Authors:** Tea Soon Park, Ludovic Zimmerlin, Rebecca Evans-Moses, Justin Thomas, Jeffrey S. Huo, Riya Kanherkar, Alice He, Nensi Ruzgar, Rhonda Grebe, Imran Bhutto, Gerard Lutty, Elias T. Zambidis

**Author notes:** Correspondence: Elias T. Zambidis, MD, PhD, Institute for Cell Engineering, and Sidney Kimmel Comprehensive Cancer Center, The Johns Hopkins University School of Medicine 733 N. Broadway, MRB 755, Baltimore, MD 21205.

## Abstract

Vascular regenerative therapies with conventional human induced pluripotent stem cells (hiPSC) currently remain limited by high interline variability of differentiation and poor efficiency for generating functionally transplantable vascular progenitors (VP). Here, we report the advantage of tankyrase inhibitor-regulated naïve hiPSC (N-hiPSC) for significantly improving vascular cell therapies. Conventional hiPSC reprogrammed from type-1 diabetic donor fibroblasts (DhiPSC) were stably reverted to naïve epiblast-like state with high functional pluripotency with a cocktail of LIF and three small molecules inhibiting the tankyrase, MEK, and GSK3β signaling pathways (LIF-3i). Naïve diabetic VP (N-DVP) differentiated from naïve DhiPSC (N-DhiPSC) expanded more efficiently, possessed higher proliferation, possessed more stable genomic integrity and displayed higher *in vitro* vascular functionality than primed diabetic VP (DVP) generated from isogenic conventional DhiPSC. Moreover, N-DVP survived, migrated, and engrafted *in vivo* into the deep vasculature of the neural retinal layers with significantly higher efficiencies than isogenic primed DVP in a murine model of ischemic retinopathy. Epigenetic analyses of CpG DNA methylation and histone configurations at developmental promoters of N-hiPSC revealed tight regulation of lineage-specific gene expression and a de-repressed naïve epiblast-like epigenetic state that was highly poised for multi-lineage transcriptional activation. We propose that reprogramming of patient donor cells to a tankyrase inhibitor-regulated N-hiPSC may more effectively erase epigenetic aberrations sustained from chronic diseases such as diabetes for subsequent regenerative therapies. More broadly, tankyrase inhibitor-regulated N-hiPSC represent a new class of human stem cells with high epigenetic plasticity, improved multi-lineage functionality, and potentially high impact for regenerative medicine.

## INTRODUCTION

The human retina is dependent on an intact, functional vasculature. If either the retinal or choroidal vasculature become compromised, neurons and supporting cells in ischemic areas rapidly die. During progressive diabetic retinopathy (DR), ischemic death of retinal pericytes and endothelial cells (EC) ^1–4^ leads to acellular vascular segments, rapid death of retinal neurons, microglial stimulation, secondary inflammation, macular edema, and subsequent retinal damage from proliferative neovascularization ^5, 6^. If acellular retinal capillaries could be regenerated with patient-specific cellular therapies, neuronal death and pathological neovascularization could be halted or reversed. Human induced pluripotent stem cell (hiPSC) cell therapies offer a versatile patient-specific approach for *de novo* regeneration of pericytic-EC ^7, 8^. Durable, albeit limited long-term *in vivo* engraftment of conventional hiPSC-derived vascular progenitor (VP) cells into the ischemic retina was previously reported ^7^. However, despite the potential and rapid advance of ocular regenerative medicine ^9, 10^, conventional hiPSC lines currently remain limited by highly variable differentiation efficiency and poor *in vivo* functionality of VP derived from them.

One critical variable impacting the differentiation efficiency and functional pluripotency of conventional hiPSC is the developmental, biochemical, and epigenetic commonality of hiPSC with ‘primed’ murine post-implantation epiblast stem cells (mEpiSC), which possess a more restricted pluripotency than inner cell mass-derived mouse ESC (mESC) (Reviewed in ^11^). Conventional hiPSC cultures adopt a spectrum of mEpiSC-like pluripotent states with highly variable lineage-primed gene expressions and post-implantation primed epiblast epigenetic marks that result in inconsistent or diminished differentiation ^12, 13^. Moreover, epigenetic aberrations in diseased states such as diabetes further inhibit efficient donor cell reprogramming to functional pluripotent states ^14–18^.

Naïve hiPSC (N-hiPSC) with more primitive pre-implantation epiblast phenotypes, decreased lineage priming, improved epigenetic stability, and higher functionality of differentiated progenitors may solve these obstacles, but this potential has not yet been demonstrated. Several groups have reported various complex small molecule approaches that putatively captured human ‘naïve-like’ pluripotent molecular states that are more primitive than those exhibited by conventional, primed hiPSC (reviewed in ^11^). However, many of these human naïve-like states exhibited karyotypic instability, global loss of parental genomic imprinting, and impaired multi-lineage differentiation performance. In contrast, a small molecule-based naïve reversion method that employed the tankyrase inhibitor XAV939 along with leukemia inhibitory factor (LIF), and the classic 2i cocktail of GSK3β and MEK/ERK inhibition (LIF-3i) avoided these caveats ^12, 13^. The LIF-3i method rapidly reverted conventional, lineage-primed human pluripotent stem cells (hPSC) to a stable pluripotent state that adopted biochemical, transcriptional, and epigenetic features of the human pre-implantation naïve epiblast. LIF-3i-reverted N-hPSC possessed greater differentiation potency than conventional hiPSC, and this improved functional pluripotency was directly potentiated by inclusion of the small molecule tankyrase/PARP (poly ADP ribose polymerase) inhibitor XAV939 to the classical 2i cocktail. Interestingly, inclusion of XAV939 into various other stem cell growth media has now further identified a new class of pluripotent stem cells with improved functionalities and ‘expanded pluripotency’ ^19–21^. For example, inclusion of XAV939 expanded the functionality of murine naïve epiblast-like PSC ^19, 20^, as well as hESC with a primed epiblast phenotype ^20, 21^ for generating both embryonic and extra-embryonic derivatives ^19, 21^.

Here, we demonstrate for the first time that the epigenetic obstacle of lineage priming and high interline variability of vascular lineage differentiation from normal and diseased conventional hiPSC can be eliminated by reversion to this novel tankyrase inhibitor-regulated naïve pluripotent state. Notably, naive diabetic VP (N-DVP) differentiated from patient-specific naïve diabetic hiPSC (N-DhiPSC) maintained greater genomic stability, higher expression of vascular identity markers, decreased non-lineage gene expression, and were superior in migrating to and re-vascularizing the deep neural layers of the ischemic retina than conventional diabetic VP generated from the same genotypic-identical isogenic DhiPSC line. We propose that naïve VP (N-VP) will have great potential for treatment of vascular ischemic disorders. Moreover, reprogramming of patient donor cells to a tankyrase inhibitor-regulated naïve human pluripotency may have wide impact in regenerative medicine by more effectively erasing dysfunctional epigenetic memory sustained from chronic diseased states such as diabetes.

## RESULTS

### LIF-3i naïve reversion of conventional, primed hiPSC lines significantly improved their multi-lineage differentiation potency

Culture of conventional hiPSC with a small molecule cocktail of LIF, the tankyrase inhibitor XAV939, the GSK3β inhibitor CHIR99021, and the MEK inhibitor PD0325901 (LIF-3i) conferred a broad repertoire of normal, non-diseased hiPSC ^12, 13^ (**Table S1**) with molecular and biochemical characteristics that are unique to naïve pluripotency, including increased phosphorylated STAT3 signaling, decreased ERK phosphorylation, global 5-methylcytosine CpG hypomethylation, genome-wide CpG demethylation at ESC-specific gene promoters, and dominant distal OCT4 enhancer usage ^12, 13^. LIF-3i-reverted N-hiPSC maintained normal karyotypes (**Table S2**), and were devoid of systematic loss of imprinted CpG patterns or irreversible demethylation defects reported in other naïve reversion systems, and that were attributed to prolonged culture with MEK inhibitors ^22–24^.

LIF-3i reversion of a broad repertoire of non-diseased conventional, primed hiPSC and hESC was reported to decreased lineage-primed gene expression, and diminished the interline variability of directed differentiation typically observed amongst independent primed, conventional hPSC lines ^12, 13^. For further validation, a cohort of isogenic (genotypically-identical) naïve vs. primed, conventional normal (non-diabetic) cord blood (CB)- and fibroblast-derived hiPSC and hESC lines (**Table S1**) were differentiated in parallel using established multi-lineage differentiation protocols (**Fig. S1;** see **Methods**). In contrast to reports of difficult directed differentiations of human naïve states, or requirement for culture back to a primed pluripotent state ^25, 26^, LIF-3i-reverted N-hiPSC differentiated *directly* to all three germ layer lineages with significantly higher efficiencies and more rapid kinetics of differentiation than their isogenic primed, conventional hiPSC counterparts. This high multi-lineage differentiation potency of N-hPSC did not require an additional ‘re-priming’ culture step. For example, in identical neural induction medium conditions, N-hiPSC produced significantly higher levels of Nestin^+^SOX1^+^ neural progenitors than their isogenic primed hiPSC counterparts (**Fig. S1a**). Additionally, neural ectodermal progenitor cells from N-hiPSC more efficiently differentiated into elongated neurites expressing Tuj1 than their isogenic primed hiPSC counterparts (**Fig. S1b**). Similarly, LIF-3i N-hiPSC vs their isogenic primed hiPSC counterparts were directly differentiated in parallel cultures into definitive endoderm; N-hiPSC generated significantly higher quantities of FOXA2^+^ progenitors at levels surpassing their isogenic primed hiPSC controls (**Fig. S1c**). Finally, LIF-3i-reverted N-hiPSC and hESC differentiated with significantly higher efficiencies than their isogenic primed hESC and hiPSC controls into vascular mesoderm (*e.g.,* CD31^+^CD146^+^, KDR^+^, CD140b^+^, CD34^+^, and CD143^+^ VP), and regardless of hiPSC genotypic background or donor source (*i.e*., fibroblast or cord blood derived; *n*=5) (**Fig. S1d,e**). As previously reported ^7, 12, 34, 35^, the improved differentiation performance of N-hiPSC erased the poor efficiency and interline variability of vascular lineage differentiation routinely observed between independent conventional fibroblast-derived hiPSC lines.

To further validate the functional pluripotency of normal (non-diabetic) fibroblast-derived N-hiPSC *in vivo*, we performed multi-lineage differentiation performance in teratoma assays (Fig. 1). Gross histological analysis of eight-week old teratomas from primed and naïve normal (non-diabetic) fibroblast-hiPSC demonstrated that although both robustly generated lineages of all three germ layers, there were marked quantitative differences between isogenic primed and naive-reverted fibroblast-hiPSC in generating teratoma organoid structures. Whereas primed fibroblast-hiPSC produced well-formed cystic teratomas with strong bias toward mesodermal cartilage differentiation (Fig. 1b), N-hiPSC-derived teratomas generated more homogenous and robust distribution of multiple structures from all three germ layer lineages, and with significantly greater number per cross section of endodermal (gut) structures, neuro-ectodermal (neural rosettes, retinal pigmented epithelial) structures, and mesodermal (cartilage) structures (Fig. 1a,b). Furthermore, fibroblast-derived N-hiPSC teratomas (*n*=3) generated proliferating multi-lineage organoid structures with significantly higher proliferative indices (*e.g*., 30-50% Ki67^+^ in CK8^+^ gut endodermal, NG2^+^ mesodermal cartilage structures) than their isogenic primed fibroblast-hiPSC counterpart (Fig. 1c,d).

**Figure 1.**
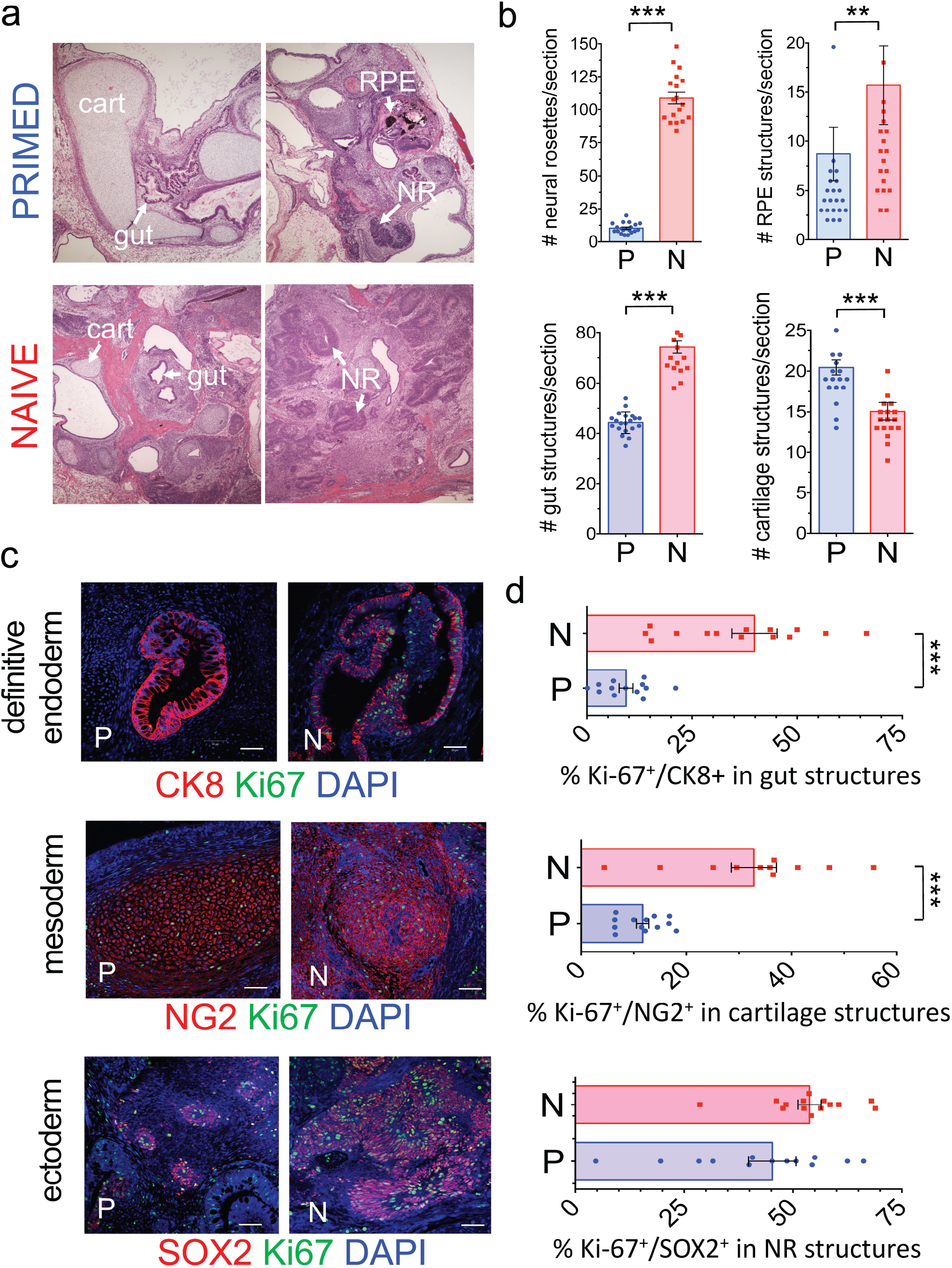
Multi-lineage teratoma organoid quantifications in isogenic primed vs. naïve non-diabetic hiPSC. The normal, non-diabetic human fibroblast-hiPSC line C1.2 (**Table S1**) was cultured in parallel in either primed, conventional E8 (PRIMED; P) or LIF-3i/MEF naïve (NAÏVE; N) conditions prior to parallel injections into sibling NOG mice (5×10^6^ cells/site) for teratoma assays. Paraffin sections of 8-week-old N vs P teratomas were evaluated and individual microscopic teratoma sections quantified by (**a, b**) H&E staining, or (**c, d**) immunofluorescence (IF) staining. Shown are individual tissue section measurements from at least 3 independent teratoma experiments quantified for organoid structures and markers of endodermal (Cytokeratin 8^+^ (CK8); gut/glandular structures), mesodermal (NG2+ chondroblasts), and ectodermal (SOX2^+^ neural rosettes) lineages along with the proliferation marker Ki-67, as described in Methods. Scale Bar = 50 μm. ** = *p* < 0.01; *** = *p* < 0.001 (Mann-Whitney tests).

### Reprogramming of skin fibroblasts of a type-1 diabetic donor to conventional DhiPSC and subsequent naïve reversion to N-DhiPSC

To test the therapeutic potential of embryonic VP derived from vascular disease-affected fibroblast-hiPSC, we generated several independent conventional SSEA4^+^TRA-1-81^+^ DhiPSC lines from type-1 diabetic donor skin fibroblasts using a modified version of a non-integrative 7-factor episomal reprogramming system ^27–29^ (**Fig. S2a-c**). Reversion of conventional, primed normal non-diabetic hiPSC or DhiPSC with the LIF-3i naïve culture system that included the tankyrase/PARP inhibitor XAV939 (**Fig. S2d**), resulted in changes from flattened (**Fig. S2b**) to dome-shaped SSEA4^+^TRA-1-81^+^ N-DhiPSC colony morphologies (**Fig. S2e**). The primed to naive transition was accompanied by activation of protein expressions of a panoply of naïve-specific pluripotency factors (Fig 2a) (*e.g*., NANOG, KLF2, NR5A2, TFCP2L1, STELLA/DPPA3, and E-CADHERIN), as well as naïve ESC-specific proteins, that included phosphorylated STAT3 and TFAP2C ^31^ (Fig 2b,c). All N-DhiPSC lines possessed normal karyotypes (**Table S2, Fig. S2e**), and generated robust tri-lineage teratomas with significantly higher differentiation performances of organoid structures representing all three germ layers than their primed isogenic counterparts (Fig. 2d**, S2f**), and with comparable efficiencies to non-diabetic N-hiPSC (Fig. 1).

**Figure 2.**
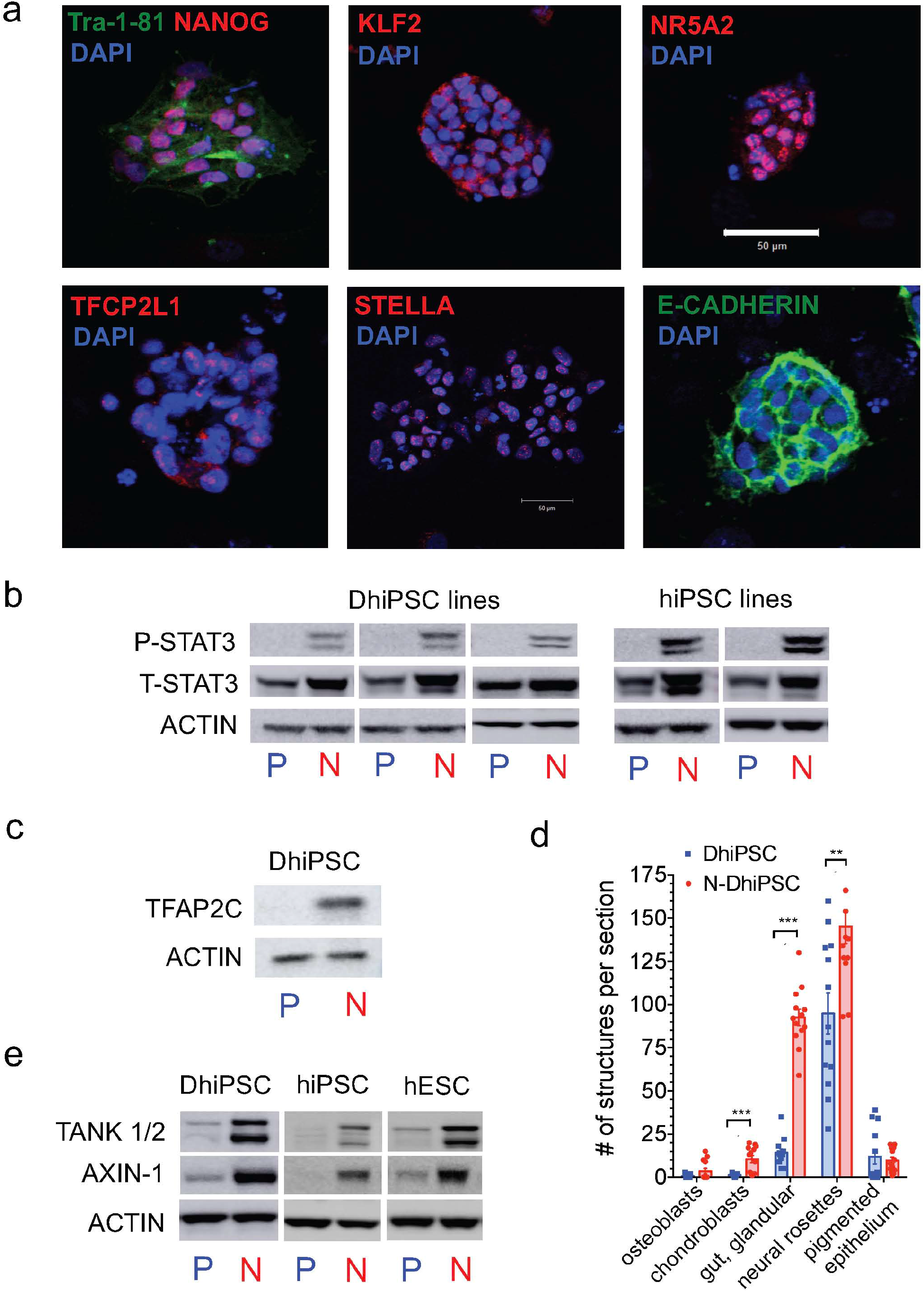
Generation of isogenic conventional and naive DhiPSC lines. (**a**) Immunofluorescent stains of N-DhiPSC (line E1C1) following LIF-3i reversion from its conventional primed state for general pluripotency factors (TRA-1-81, NANOG, OCT4) and naïve pluripotency proteins (KLF2, NR5A2, TFCP2L1, STELLA/DPPA3, E-CADHERIN; Scale Bar = 50 μm). Primed- *vs*. naïve-cultured isogenic hPSC line protein lysates were prepared from 3 independent DhiPSC lines (E1C1, E1CA1, E1CA2), an hESC line (H9), and normal non-diabetic fibroblast-hiPSC lines (C1.2, C2). Western blots were performed of primed (P) vs naïve (N) lysates of isogenic DhiPSC line E1C1, normal CB-iPSC line E5C3, and hESC line H9, with ACTIN or total STAT3 serving as internal loading controls for each blot. (**b**) expressions of phosphorylated (P-STAT3) and total STAT3 (T-STAT3; control) in isogenic primed (P) vs naive (N) conditions from (upper panel) 3 independent DhiPSC lines (E1C1, E1CA1, E1CA2, and (lower panel) 2 independent, isogenic normal non-diabetic fibroblast-hiPSC lines (C1.2, C2) in primed vs naïve conditions. (**c**) naïve-specific expression of pluripotency factor TFAP2C ^31^ is shown for DhiPSC line E1C1 in primed (P) vs naïve (N) isogenic conditions (**d**) Isogenic teratoma organoid quantifications from diabetic hiPSC line E1C1 cultured in primed (blue bar), vs naive (red bar) conditions. Shown are quantifications per cross section of mesodermal (NG2^+^ chondroblast), definitive endodermal (CK8^+^ gut/glandular cells), and ectodermal (SOX2+ neural rosettes; retinal pigmented epithelium) structures from H&E stained slides. ** = p< 0.01; *** = p < 0.001 (Mann-Whitney tests). (**e**) Western blot analysis of XAV939-inhibited proteolysis of tankyrases 1 and 2 (TANK ½) and AXIN-1 proteins from DhiPSC line E1C1, CB-hiPSC line E5C3, and hESC line H9, in isogenic naïve (N) *vs*. primed (P) conditions.

To validate the effects of XAV939 inhibition of tankyrase-PARP activity in DhiPSC, we verified proteolytic inhibition of key proteins targeted by tankyrase PARylation, including AXIN1 (which synergizes with the GSK3β inhibitor to stabilize the activated **β-**catenin complex ^20^), and tankyrase 1 (PARP-5a) and tankyrase 2 (PARP-5b) proteins (which self-regulate their own proteolysis by auto-PARylation) ^30^. Accordingly, chemical inhibition of their degradation resulted in high accumulated levels of tankyrases 1/2 and AXIN1 in LIF-3i-reverted N-DhiPSC that was comparable to non-diabetic fibroblast- and non-diabetic cord blood (CB)-derived hiPSC lines (Fig. 2e).

### LIF-3i naïve reversion improved the efficiency of vascular lineage differentiation of diabetic donor-derived conventional hiPSC lines

We previously identified a FACS-purified CD31^+^CD146^+^CXCR4^+^ embryonic VP population differentiated from conventional (non-diabetic) hiPSC that possessed both endothelial and pericytic functionalities ^7^. We demonstrated that Ischemia-damaged retinal vasculature could be repaired by transplantation of these hiPSC-derived CD31^+^CD146^+^ embryonic VP generated from conventional hiPSC, and that they possessed prolific endothelial-pericytic differentiation potential and engrafted and rescued degenerated retinal vasculature following ocular ischemia-reperfusion (I/R) injury.

We employed an optimized isogenic primed vs naïve hiPSC version of our original VP differentiation system ^7^ (Fig. 3a), and similar to results obtained for non-diabetic hiPSC ^32, 33^ (**Fig. S1d,e**), we found that CD31^+^CD146^+^ DVP cells were more efficiently generated from N-DhiPSC, and with more rapid differentiation kinetics than their primed, conventional isogenic DhiPSC counterparts (Fig. 3b,c**; Fig. S3a,b**). Kinetic analyses of parallel vascular differentiation cultures of N-DhiPSC vs primed isogenic DhiPSC revealed higher expressions of vascular surface markers (*e.g*., CD34, CD31, CD146, CD144), and a more rapid decrease of pluripotency markers (*e.g.,* SSEA4, TRA-1-81) in these naïve diabetic VP (N-DVP). Naive reversion permitted comparable efficiencies of generation of CD31^+^CD146^+^CXCR4^+^ VP populations *regardless of conventional hiPSC donor source* (*i.e*., diabetic or non-diabetic). For example, naïve-reverted fibroblast-derived N-DhiPSC and non-diabetic cord blood (CB)-derived N-CB-hiPSC lines both generated similar efficiencies of VP, despite a previously reported higher VP differentiation efficiency of conventional CB-hiPSC compared to conventional fibroblast-derived hiPSC ^7^ (Fig. 3c). Following MACS enrichment of CD31^+^CD146^+^ VP populations from vascular differentiation cultures of both non-diabetic and diabetic hiPSC, naïve and primed VP populations both displayed similar and equalized expressions of a broad array of vascular markers (*e.g*., CD31, CD34, CD144 and CD146) (**Fig. S3c**). Moreover, endothelial progenitor-specific (CD31^+^CD105^hi^CD144^+^), pericytic (CD31^+^CD90^+^CD146^+^) populations ^36^ (**Fig. S3c**), and subcellular endothelial-specific organelles (*e.g.,* Weibel-Palade (WP) bodies, and coated pits; Fig. 3d;) were similarly expressed in both DVP and N-DVP; albeit with some minor differences (*e.g*., more abundant transcytotic endothelial channels (TEC) (Fig. 3d).

**Figure 3.**
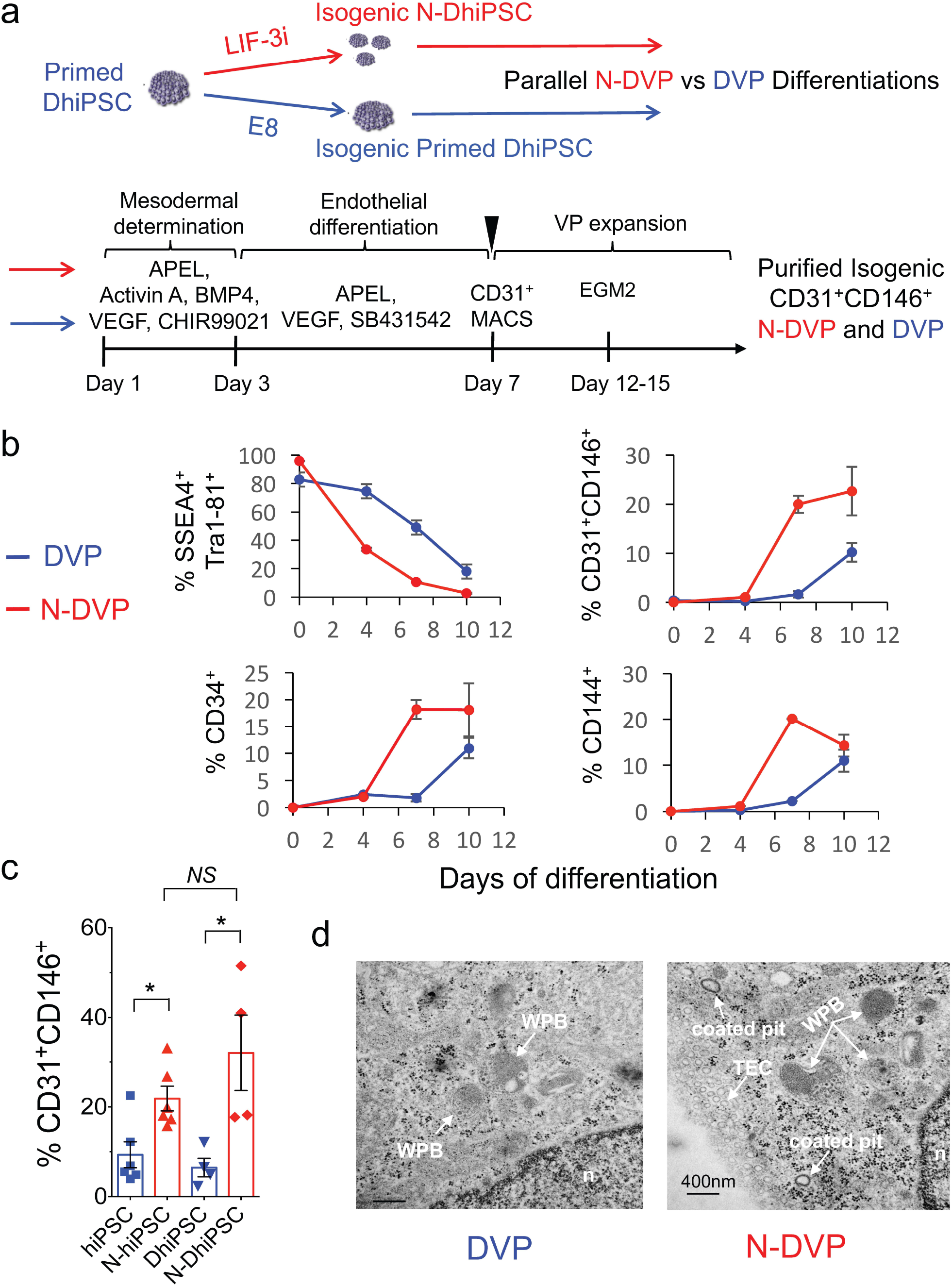
Vascular differentiations of primed vs. naïve non-diabetic and diabetic hiPSC. (**a**) (Upper panel) schematic of experimental design of isogenic primed (E8) vs naïve (LIF-3i) sibling DhiPSC, differentiated in parallel following 5-7 passages in their respective primed vs naïve culture conditions, as described in Methods. (Lower panel) defined xeno-free APEL vascular differentiation system. Shown are indicated growth factors and inhibitor molecules, as described in Methods. Day 7 differentiation cultures were enriched for CD31-expressing VP using magnetic-activated cell sorting (MACS); CD31^+^ VP co-express CD146 post-MACS enrichment. These CD31^+^CD146^+^ VP populations were further expanded for several passages in EGM2 medium prior to *in vitro* characterization or injection into I/R-injured murine NOG eyes. (**b**) Kinetics of surface protein expressions by flow cytometry during APEL vascular differentiation for pluripotency markers (SSEA4, TRA-1-81) and vascular markers (CD31, CD146, CD34, CD144) from isogenic primed (blue) and naïve (red) DhiPSC lines. Shown are mean results with SEM of two independent differentiation experiments of N-DhiPSC and their isogenic primed DhiPSC counterparts (lines E1C1, E1CA1; *n*=2). (**c**) Average percentages of CD31^+^CD146^+^ cells obtained from APEL differentiation cultures of isogenic independent non-diabetic CB-iPSC and diabetic hiPSC lines on differentiation days 7-8 (*i.e*., day of CD31+ sorting (see scheme above). Results of independent experiments are shown for differentiation of the non-diabetic CB-iPSC line E5C3 and two DhiPSC lines (E1CA1, E1CA2) starting from simultaneous and isogenic primed and naïve cultures. * = p < 0.05 (unpaired two-tailed t tests). (**d**) Transmission electron microscopy (TEM) images of primed DVP and N-DVP differentiated and expanded as described above from parallel primed and naïve isogenic conditions of the DhiPSC line E1C1. WPB: Weibel-Palade body, n: nucleus, TEC: transcytotic endothelial channel; Scale Bar = 400nm.

### N-DVP possessed improved vascular functionality, lower culture senescence, and reduced sensitivity to DNA damage

Endothelium dysfunction in diabetics is characterized by poor EC survival, function, and DNA damage response (DDR). Although regenerative replacement of diseased vasculature requires high functioning cell therapies, previous studies of vascular differentiation with conventional fibroblast-hiPSC revealed poor and variable growth and expansion of vascular lineage cells, with high rates of apoptosis and early senescence ^34, 35^. To evaluate endothelial functionality of naïve CD31^+^CD146^+^ VP, purified primed DVP vs N-DVP populations were re-cultured and expanded in endothelial growth medium (EGM2). N-DhiPSC-derived N-DVP were compared to isogenic primed DhiPSC-derived DVP for *in vitro* endothelial functionality with acetylated-Dil-LDL (Ac-Dil-Ac-LDL) uptake assays (Fig. 4a), 5-ethynyl-2-deoxyuridine (EdU) cell cycle proliferation assays, (**Figs. 4b, S4b**), culture senescence (**Figs. 4c, S4a**), and quantitative Matrigel vascular tube formation (Figs. 4d**, S4c**). These studies collectively revealed that CD31^+^CD146^+^-enriched N-DVP maintained higher endothelial Ac-Dil-Ac-LDL uptake function, more enhanced proliferation, and significantly less culture senescence than isogenic primed DVP counterparts in endothelial re-culture and expansion conditions.

**Figure 4.**
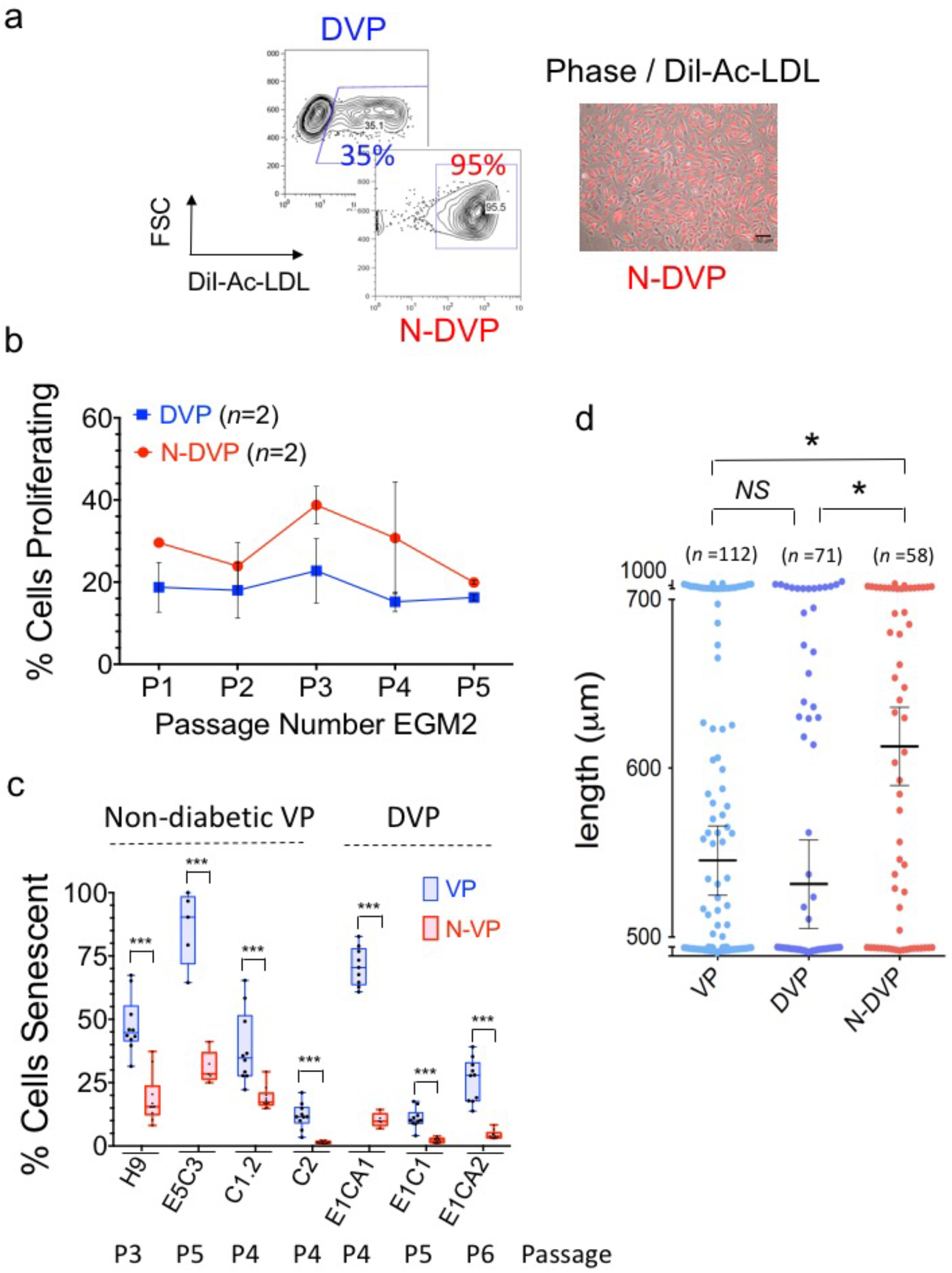
Characterization of DVP generated from primed *vs*. naïve DhiPSC. (**a**) Endothelial functionality. Shown are representative flow cytometry (left panel) and immunofluorescent Dil-acetylated-LDL (Dil-Ac-LDL) endothelial uptake assays (right panel); merged phase contrast/ Ac-Dil-LDL-labeled primed DVP vs. N-DVP cells; Scale Bar = 100μm. DVP cells were generated from primed vs naïve isogenic DhiPSC line E1CA2. (**b**) EdU proliferation assays of purified DVP after 4 passages in EGM2 post-CD31^+^ purification. N-DVP demonstrated higher proliferative capacity up to passage 3 compared to isogenic primed cells. (**c**) Expanded VP and N-VP were quantitated for senescent cells by β-galactosidase activity colorimetric assay. Shown are isogenic independent comparisons of both non-diabetic primed VP and N-VP (i.e., generated from H9 hESC, C1.2, C2 fibroblast-hiPSC lines) and diabetic DVP and N-DVP (*i.e*., generated from E1CA1, E1CA2, E1C1 DhiPSC lines). Each quantitation is an independent measurement of EGM2 cultures at indicated matched passages for each VP and N-VP type. *** = p < 0.001 (multiple unpaired t tests). (**d**) Quantification of vascular tube lengths formed from *in vitro* Matrigel tube assays from primed normal VP (E5C3) and isogenic primed DVP vs N-DVP (line E1CA2).

To further evaluate the relative resistance of N-DVP to senescence, we probed genomic integrity maintenance by assaying for sensitivity to double stranded DNA breaks (DSBs) following treatment with the radiation damage mimetic neocarzinostatin (NCS), which triggers both DDR and pH2AX-mediated reactive oxygen species (ROS) signals ^37^. Expression of phosphorylated p53 protein (P-p53), phosphorylated H2AX (pH2AX), RAD51, RAD54, phosphorylated DNA-PK (pDNA-PK), which are all normally activated briefly following DDR and mediate repair of DSBs, were compared in re-cultured and expanded primed DVP vs N-DVP, before and after treatment with NCS (Fig. 5a-c, **Fig. S5**). These studies revealed that levels of the DDR and DSB protein network (*e.g*., pH2AX, P-p53, RAD51, RAD54, and p-DNA-PK) were all globally reduced in their expressions in both non-diabetic N-VP and N-DVP; both before *and* following NCS DNA damage exposure. Collectively, these data demonstrated that N-DVP may mediate a reduced sensitivity to stress-induced DNA damage than do primed DVP. These studies also suggested an improved genomic integrity of N-VP and N-DVP *relative to VP generated from conventional counterparts* of both normal hiPSC and diseased DhiPSC, respectively.

**Figure 5.**
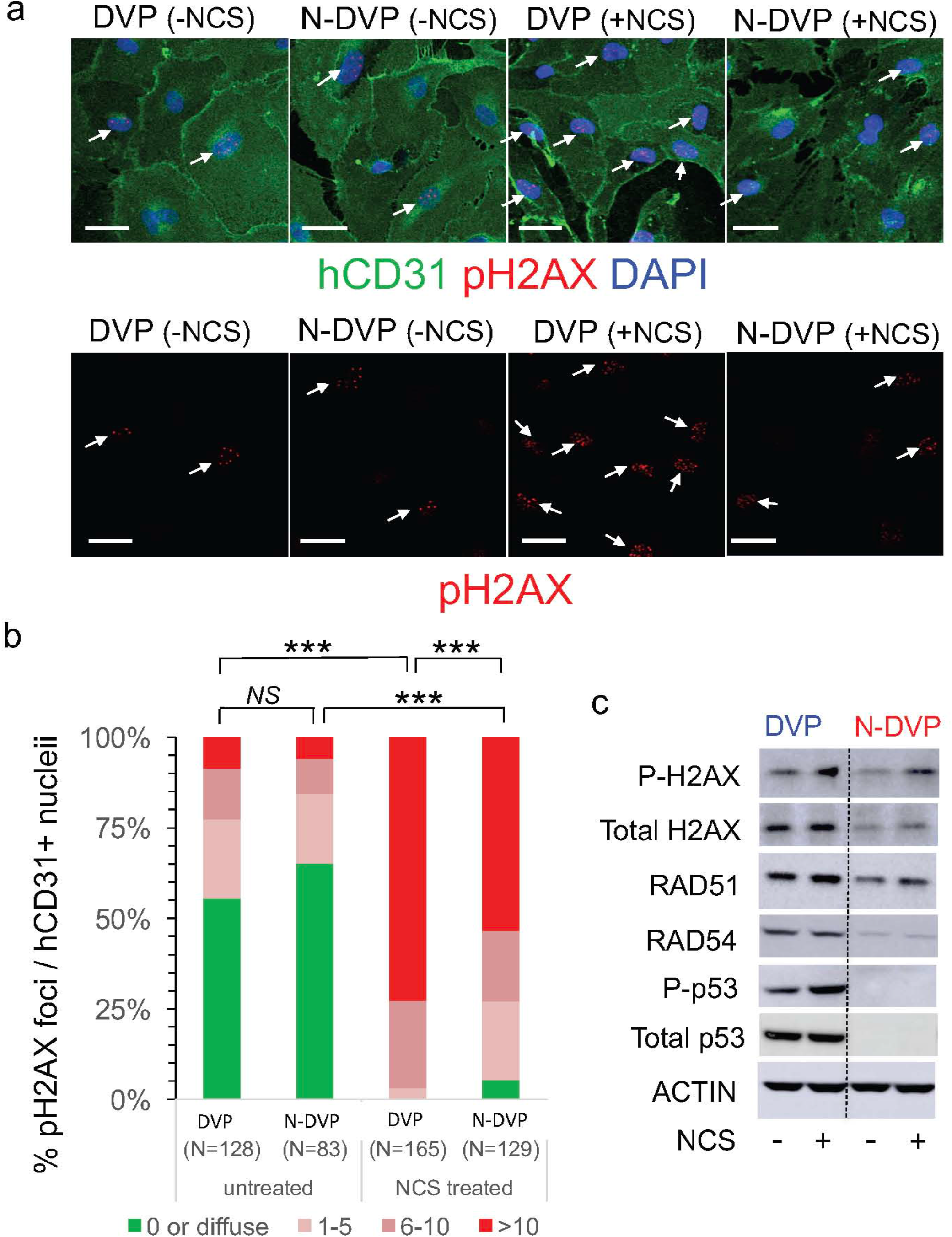
DNA damage responses in primed DVP vs N-DVP. (**a**) purified and expanded CD31^+^ DVP or N-DVP (line E1CA2) were treated with the radiomimetic drug NCS for 5 hours before fixation and staining with antibodies for detection of human CD31^+^ cells and phosphorylated H2AX (pH2AX) positive nuclear foci (*i.e*., DAPI co-staining) to reveal double-strand DNA breaks (arrows); Scale Bar = 50μm. (**b**) Quantification of pH2AX foci per nuclei in isogenic primed vs naïve DVP with or without induction of DNA damage with NCS. Shown are numbers of DAPI+ nuclei per field with no pH2AX foci (green), and DAPI+ nuclei with 1-5 foci (light pink), 6-10 (dark pink) and >10 pH2AX foci (red). *** = p<0.0001; Chi-Square tests (**c**) Lysates of primed DVP and and N-DVP (line E1C1) cultured in EGM2 and treated with and without NCS for 5 hours were analyzed by Western blotting for expressions of proteins activated by DNA damage and apoptosis *(i.e*., phosphorylated H2AX (P-H2AX), RAD51, RAD54, and phosphorylated p53 (P-p53).

### N-DVP injected into the vitreous of eyes survived, migrated into the neural retina, and engrafted into ischemia-damaged retinal vasculature with high efficiency

To evaluate the potential of N-DVP for *in vivo* engraftment and repair of ischemic retinal vessels, we employed our previously described humanized experimental NOG mouse model of ocular ischemia-reperfusion (I/R) injury that allows the engraftment of human VP in an *in vivo* ischemic retinal niche (Fig. 6a) ^7^. This model has been utilized extensively in rodents to recreate the ischemic damage observed in diabetic retinopathy ^2^. Intraocular pressure can be experimentally elevated to 90 mm Hg for 90 minutes in mice, followed by allowance of vascular reperfusion. This manipulation results in loss of retinal vasculature and apoptotic death of ischemic retinal neurons ∼7 days following I/R injury. Although this rodent model does not exactly simulate the sequence of events in diabetic retinopathy, it *produces the same pathology seen in all ischemic retinal diseases: acellular capillaries and ischemic death of neurons*. In our humanized immune-deficient mouse model, the sequential loss of murine host retinal vasculature ECs following ocular I/R injury was detected with an antibody specific to mouse anti-CD31 (mCD31), and the vascular basement membrane was detected with a murine-specific anti-collagen type-IV (mCol-IV) antibody. This approach is feasible because despite ischemic damage to acellular capillaries, the basement membrane shared by EC and pericytes in retinal capillaries remains intact. In this model, ischemic damage is more severe in capillaries and veins presumably due to their higher collapsible nature under increased intraocular pressure compared to arteries.

**Figure 6.**
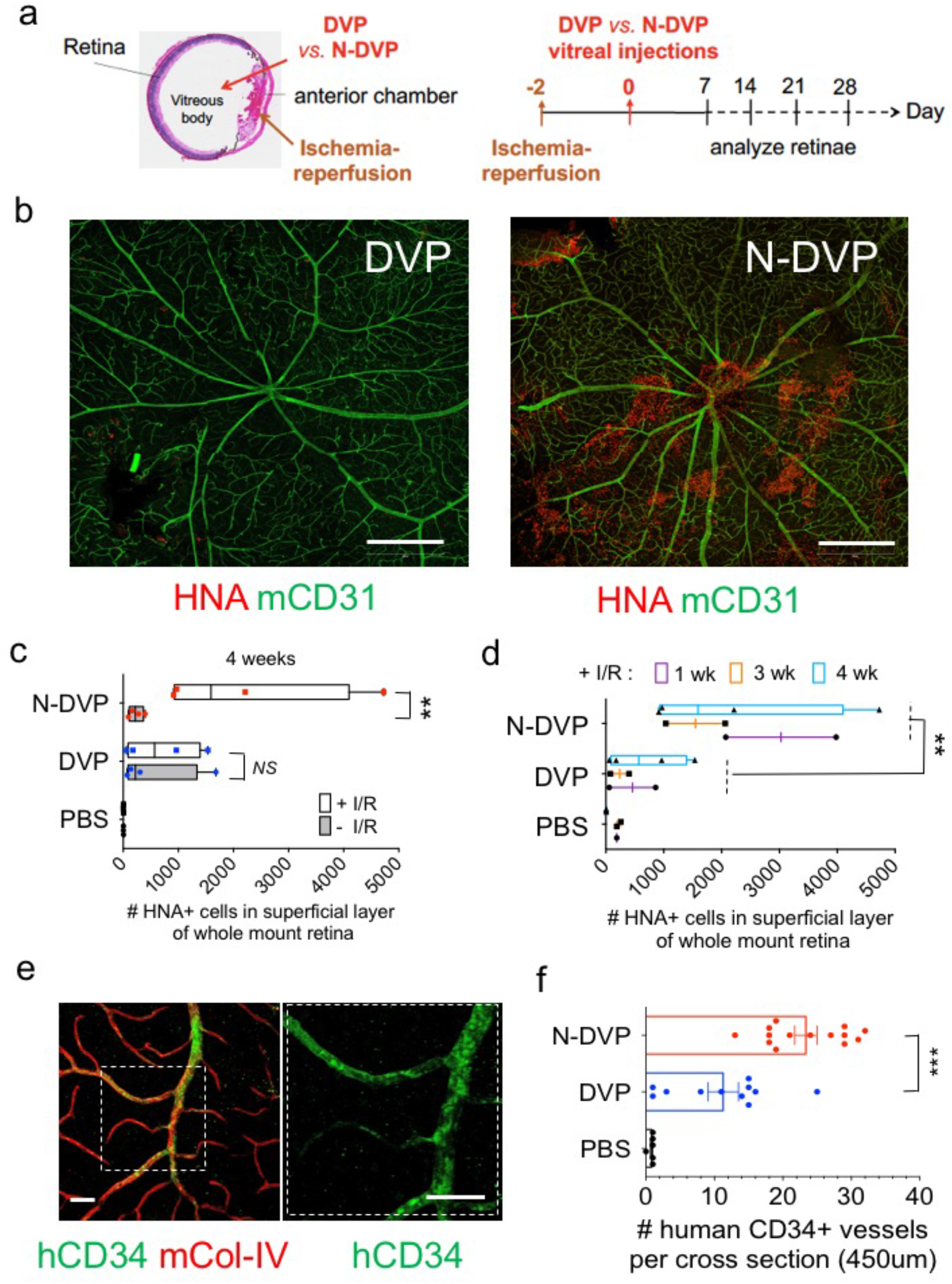
Efficient survival and human vascular engraftment of N-DVP in I/R-injured murine retinae. (**a**) Schematic of NOG mouse ocular I/R experimental system for testing *in vivo* functionality of human primed DVP vs N-DVP, modified from *Park et al*, *Circulation,* 2014**^7^**. Shown are anatomical structures where I/R (anterior chamber) and human DVP and N-DVP cell injections (vitreous body) were performed (left panel). Schematic of timeline for I/R injury surgery, human cell DVP injections (Day 0), and days of analysis of human cell survival and engraftment (right panel). (**b**) Human DVP cell survival at the superficial layer of murine retina at three weeks following injection of 50,000 DVP or N-DVP into the vitreous of I/R-treated NOG mouse eyes. Flat whole-mounted retinae were stained with antibodies for human-specific HNA (red), and tile scanned by confocal microscopic imaging (10x objective, 9 x 9 tiles). Shown are representative whole retinal images with HNA^+^ cells from primed DVP cell-injected (left panel) *vs*. N-DVP cell-injected (right panel) eyes. Scale bars = 500 μm. (**c**) Quantitation of HNA^+^ cells detected in the outer superficial layers of whole mount retinae following treatment of eyes with and without I/R, and injected with either primed DVP or N-DVP at (**c**) 4 weeks or (**d**) at 1, 3, and 4 weeks following DVP *vs*. N-DVP vs. control saline (PBS) injections, in eyes treated with and without I/R injury. Shown are the mean numbers from independent eye experiments of total HNA^+^ cells counted with imaging software per superficial layer of each whole-mounted retinae (whole field). ** = *p* < 0.01 (Mann-Whitney tests). (**e**) Human vascular engraftment. Whole-mounted retinae of I/R-injured eyes of NOG mice 2 weeks following DVP *vs.* N-DVP *vs.* PBS injections into ischemia-injured eyes were immuno-stained with human CD34 (hCD34) to detect human endothelial engraftment. Antibodies for murine collagen type-IV (mCol-IV) were also employed (to detect murine blood vessel basement membrane), and murine CD31 (mCD31) (to detect murine endothelium). (**f**) The number of CD34+ human-murine chimeric vessels per 450 μm cross-section were quantitated via confocal microscopy and imaging software. Shown are results of independent measurements. *** = *p* < 0.001 (multiple unpaired t tests).

CD31^+^CD146^+^-sorted human DVP cells were differentiated from isogenic primed vs N-DhiPSC as above, cultured briefly in EGM2, and 50,000 primed DVP or N-DVP cells were injected in parallel directly into the vitreous body of NOG recipient eyes 2 days following I/R injury (Fig. 6a). Human cell viability, intra-retinal migration, and engraftment in murine retina was evaluated at 1, 2, 3, and 4 weeks following human DVP injections with human-specific anti-human nuclear antigen (HNA) antibody staining. Analysis between 1-4 weeks following vitreous body injection in I/R-damaged eyes, HNA^+^ N-DVP were observed to survive in significantly greater frequencies than primed DVP within the superficial layer of the retina (Fig. 6b-d**, Fig. S6a**). Additionally, HNA^+^ N-DVP were observed to assume adherent abluminal pericytic and luminal endothelial positions with greater frequencies (**Fig. S6b),** and appeared to favor venous engraftment of blood vessels with large diameter than arteries suggesting a preferential migration in response to injury signals ^7^. In contrast, and as previously demonstrated for non-diabetic fibroblast-derived hiPSC ^7^, primed DVP cells examined at 2-4 weeks post-injection survived poorly and migrated inefficiently into ischemia-damaged blood vessels, and instead remained primarily in either the vitreous, or adherent to the adjacent superficial layer of the retina (Fig. 6b-d). N-DVP not only could be confirmed to significantly survive longer at 4 weeks post injection robustly in the superficial retinal ganglion cell layers (GCL) in response to injury (Fig. 6c), but also engrafted into murine retinal vessels with significantly greater human CD34^+^ grafting efficiencies than primed DVP (Fig. 6e,f**; S6c,d**).

Interestingly, further analysis of deeper retinal vessels in transverse sections of the neural retina with anti-human CD34 and anti-human CD31 antibodies confirmed significantly higher endothelial engraftment from N-DVP than from primed DVP (Fig. 7). At 2 weeks following I/R injury, human CD34^+^, and human CD31^+^ vessels were already readily detectable in N-DVP-injected retinae at significantly higher rates than primed DVP-injected retinae in the intima of murine host ischemia-damaged capillaries located in the ophthalmic artery distribution of the deep neural retinal layers. Notably, N-DVP efficiently migrated from the outer vascular layers of the GCL to form engrafted chimeric CD34^+^ and CD31^+^ human vessels in the deeper outer plexiform layers (OPL), inner nuclear layers (INL), and inner plexiform layers (IPL) of the murine neural retina (*e.g*., ∼5-20 CD34+ human-murine chimeric vessels per 450μm cross section areas; Fig. 7a-d). In contrast, injected primed DVP-derived human CD34^+^ and CD31^+^ cells were detected primarily only in the superficial vascular layers of the retinal GCL or the inner limiting membrane (ILM); without further significant migration into deeper neural layers (Fig. 7b,d). Collectively, these data demonstrated that N-DVP but not primed DVP migrated more efficiently from vitreous into the injured deep vascular neural retinal layers; suggesting that an efficient reparative injury-induced human-murine chimeric vasculogenesis had occurred following N-DVP (but not primed DVP) cellular therapy.

**Figure 7.**
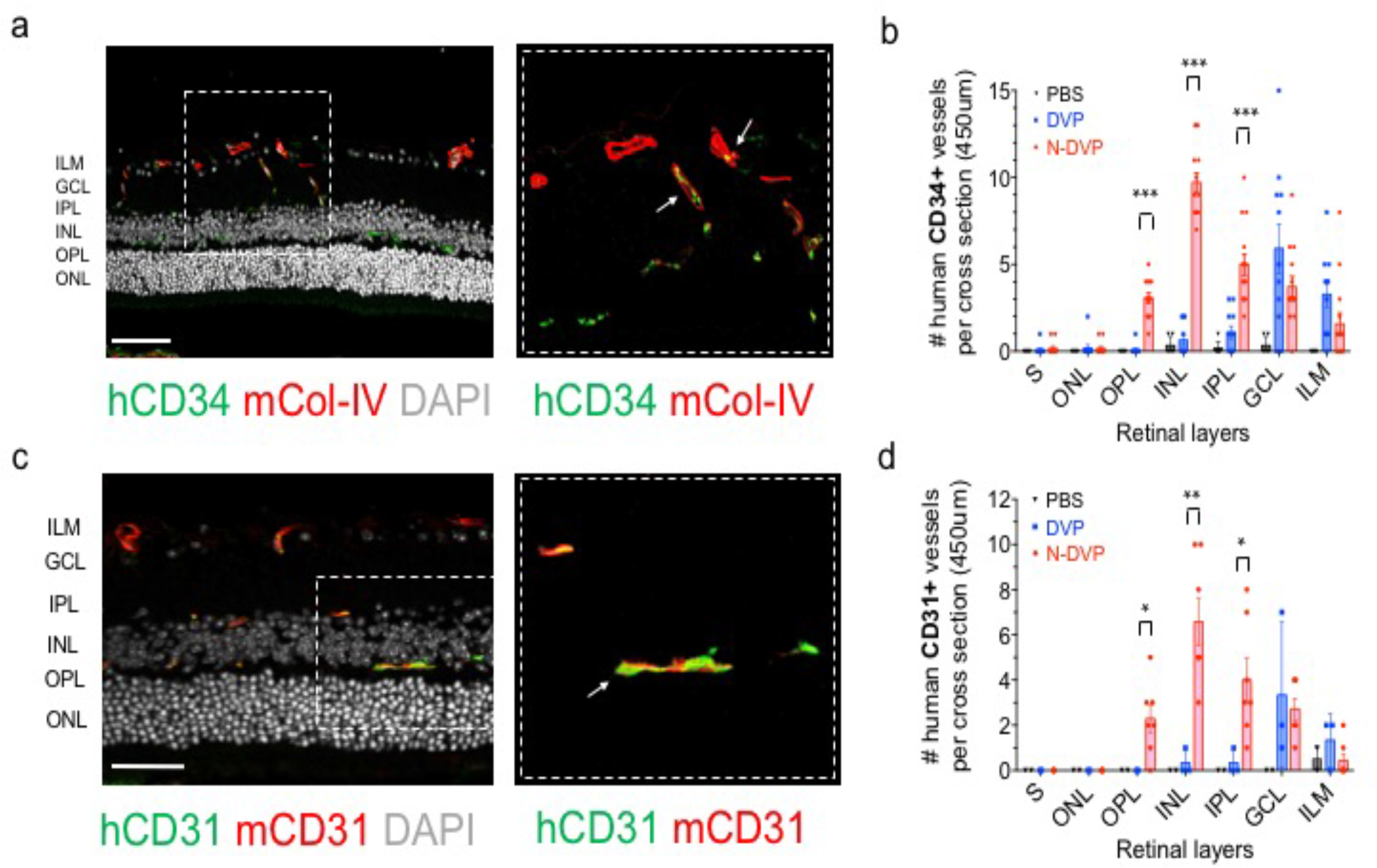
Migration of primed DVP and N-DVP into ischemia-injured blood vessels and vascular engraftment in the neural retina. Whole mount retinae of I/R-injured eyes of NOG mice were immuno-stained with either human CD34 (**a,b**; hCD34) or human CD31 (**c,d**; hCD31) antibodies to detect human endothelial cells 2 weeks following DVP *vs.* N-DVP injections. Antibodies for murine collagen type-IV (mCol-IV) were also employed to detect murine blood vessel basement membrane, and murine CD31 (mCD31) detected murine endothelium. The number of human CD34^+^ (**b**) or (**d**) human CD31^+^ cells detected within transverse layers of the murine neural retina (per 450μm retinal cross section; see **Fig. S7**) was quantitated. Each data point represents a replicate individual 450μm retinal cross section that was analyzed from I/R-treated eyes injected with saline (PBS), primed DVP or N-DVP. Human CD34^+^ or human CD31^+^ endothelial cell engraftment was enumerated in each distinct layer of neural retina shown, and demonstrated that only N-DVP migrated into the inner nuclear layer (INL) while most of the primed DVP remained primarily in the superficial ganglion cell layer (GCL). ILM: inner limiting membrane, IPL: inner plexiform layer, outer nuclear layer (ONL), OPL: outer plexiform layer, S: segments. All scale bars = 50 μm.

### N-DhiPSC were configured with de-repressed, activation-poised bivalent histone marks at key developmental promoters, and tight regulation of ‘leaky’ lineage-primed gene expression

The murine naïve pluripotent state, which has higher differentiation potential than the primed murine pluripotent state ^11^ is distinguished by chromatin poised for unbiased gene activation ^38^, global reduction of CpG DNA methylation ^39^, and decreased repressive H3K27me3 histone deposition at bivalent Polycomb repressor Complex 2 (PRC2)-regulated promoter sites ^40, 41^. To explore the molecular mechanisms that drive improved vascular functionality of LIF-3i-reverted N-DhiPSC, we next probed the epigenetic configurations that may regulate more faithful vascular gene expression in N-DVP. As was reported for non-diabetic fibroblast-N-hiPSC ^12^, all three fibroblast-N-DhiPSC lines exhibited significant reductions in global 5-methylcytosine (5MC)-associated CpG DNA methylation following LIF-3i reversion (Fig 8a). The detection of increased global levels for 5-hydroxymethylcytosine (5hMC) in N-DhiPSC relative to primed DhiPSC further suggested a potential contribution of naïve-like TET-mediated active CpG demethylation ^39^.

**Figure 8.**
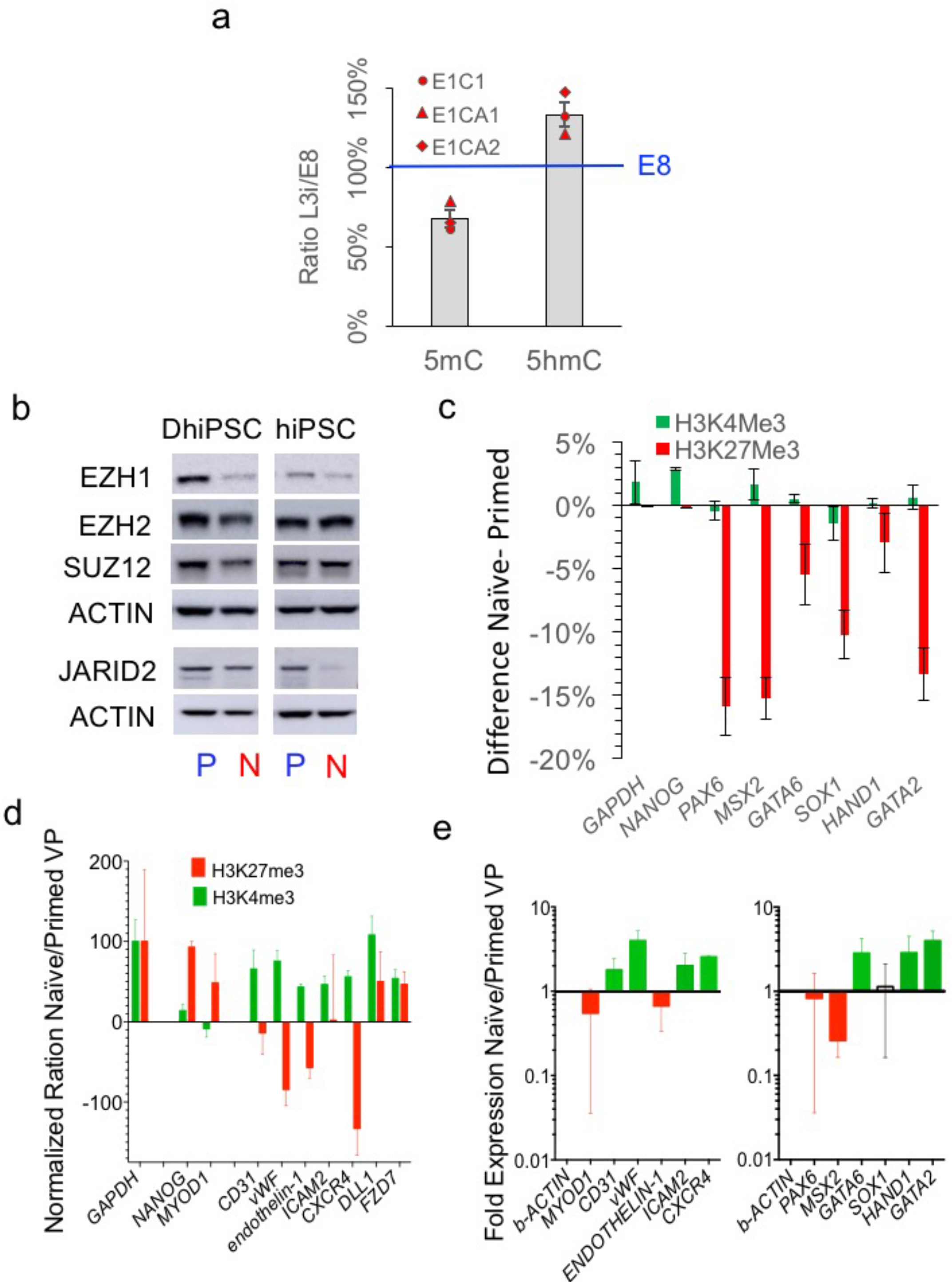
Epigenetic configurations of multi-lineage bivalent and vascular lineage-specific promoters in primed vs naïve hiPSC and VP. (**a**) Densitometric quantitation of dot immuno-blots of global levels of 5-methylcytosine (5mC) and 5-hydroxymethylcytosine (5mC) in three isogenic pairs of primed (E8) and naïve (MEF-depleted LIF-3i) DhiPSC lines. Genomic DNA samples were collected from parallel isogenic primed (E8 medium) and naïve LIF-3i) cultures. Immunoblot densities LIF-3i/E8 ratios were determined with ImageJ software at steady state conditions (200ng), and normalized at 100% for E8 values. (**b**) Western blot analysis of PRC2 components EZH1, EZH2, SUZ12, and JARID2) in primed vs naïve normal and diabetic hiPSC lysates; E1C1 fibroblast-DhiPSC line; C1.2 normal donor fibroblast-hiPSC line. (**c**) ChIP-qPCR for H3K27me3 and H3K4me3 histone marks at key known bivalent developmental promoters in primed vs naïve DhiPSC (*e.g*., PAX6, MSX2, GATA6, SOX1, HAND1, GATA2). Levels of GAPDH and NANOG are controls for actively transcribed genes. Data is presented as *differences* in percent input materials of naïve minus primed genomic DNA samples for DhiPSC line E1C1 (see **Fig S8c**). Bars represent the SEM of replicates. (**d**) ChIP-qPCR for H3K27me3 and H3K4me3 histone marks at key vascular developmental promoters in primed vs naïve VP genomic samples. Data is presented as GAPDH-normalized ratios of percent input materials between naïve and primed VP differentiated from the DhiPSC line E1C1. Results are shown as ratios of expression of isogenic N-DVP *vs*. DVP for GATA2-regulated genes (CD31, vWF, endothelin-1, ICAM2) and genes regulated by histone marks that are known to effect vascular functionality (CXCR4, DLL1, FZD7). The histone profile for GAPDH is a ‘housekeeping control gene. NANOG and MYOD1 are control gene promoters that become repressed during vascular differentiation. (**e**) qRT-PCR gene expression analysis of vascular lineage genes (left panel) and PRC2-regulated bivalent lineage-specific genes (right panel) in DVP vs. N-DVP that were differentiated from isogenic pairs of naive vs primed D-hiPSC lines (*n*=3; lines E1C1, E1CA1, E1CA2). Fold changes are normalized to beta-actin expression. All PCR primers are listed in **Table S3**.

To better elucidate the epigenetic mechanisms regulating decreased lineage priming, a simultaneous bioinformatics analyses of both microarray expression and Infinium CpG methylation arrays of lineage-specifying PRC2 gene targets (before and after LIF-3i-reversion of a broad array of isogenic hiPSC lines), revealed that a broad rewiring of the lineage-specifying machinery. CpG DNA hypomethylation at promoter sites of PRC2 genes were broadly hypomethylated in N-hiPSC lines relative to their primed counterparts with a significant decrease of expression in many lineage-specifying PRC2 targets (**Fig. S8a,b**). These studies revealed that in comparison to primed fibroblast-hiPSC, isogenic N-hiPSC displayed significantly less baseline epigenetic CpG methylation-regulated repression of lineage-specifying PRC2 genes, *despite* a *broad* silencing of lineage-primed transcriptional targets of PRC2 as previously reported ^12^ (**Fig S8c, Table S3**).

A critical mechanism for protecting naïve mouse ESC from lineage priming is via regulating the poised silencing or activation of lineage-specifying genes at bivalent H3K27me3 repressive and H3K4 activation histone marks, and RNA Polymerase II (POLII) pausing at promoter sites ^40–42^. Thus, we next assessed the protein abundance of PRC2 components which mediate repressive H3K27me3 deposition on bivalent promoters in naïve versus primed normal and DhiPSCs (Fig. 8b). Interestingly, these studies revealed significantly *decreased* abundance of multiple components of the PRC2 complex in both diabetic and non-diabetic N-hiPSC, including the enzymatic subunits EZH1, EZH2, and the cofactor subunit JARID2 ^43^ (which drives the localized recruitment of PRC2 at developmental promoters in mouse ESC). To functionally validate the activity of PRC2 targets, we employed chromatin immunoprecipitation followed by qPCR (ChIP-PCR) on previously characterized lineage-specific bivalent gene promoters (*e.g*., *PAX6, MSX2, GATA6, SOX1, HAND1, GATA2;* (Fig. 8c**, S8d, Table S3**) to investigate the levels of bivalent active (H3K4me3) and repressive (H3K27me3) histone marks at these key lineage-specifying promoters. These studies revealed significant H3K27me3 reductions (5-15% from isogenic primed E1C1 and E1CA1 DhiPSC lines) following LIF-3i reversion.

Collectively, these CpG DNA methylation and histone mark studies revealed a relatively de-repressed naive epigenetic state in N-hiPSC that appeared more poised for activation than primed DhiPSC; with a potentially decreased barrier for multi-lineage gene activation relative to primed DhiPSC. Thus, as was previously demonstrated for naïve murine ESC ^38, 40^, despite a tighter regulation of ‘leaky’ lineage-primed gene expression that was presumptively silenced through alternate naïve-like epigenetic mechanisms of bivalent promoter repression (*e.g*., promoter site RNA POLII pausing ^40^), N-hiPSC appeared poised with a lower epigenetic barrier for unbiased multi-lineage differentiation.

### N-DVP possessed vascular lineage epigenetic de-repression and reduced non-vascular lineage-primed gene expression

To determine the downstream impact of an epigenetic state with an apparently lower barrier for vascular lineage activation, we investigated the epigenetic configurations of vascular-lineage specific gene promoters in differentiated DVP by ChIP-PCR. We selected the promoters of downstream genes regulated by the PRC2-regulated factor GATA2, which promotes expression of genes of endothelial-specific identity and function (*e.g*., *CD31*, *vWF*, *endothelin-1*, and *ICAM2*) ^10^. We also selected promoters of genes known to be activated by chemical EZH2 and histone deacetylases (HDAC) de-repression in human endothelial progenitor cells (EPC) (*e.g*., *CXCR4*, *DLL1*, and *FZD7*) ^44^. CD31^+^CD146^+^ DVPs vs N-DVP were MACS-purified, briefly expanded in EGM2, and ChIP-PCR was performed on promoter sites of these genes. Strikingly, relative to primed DVP, N-DVP displayed significantly increased marks for epigenetic activation (H3K4me3) and simultaneously reduced marks of promoter repression (H3K27me3) (Fig. 8d) for genes determining vascular functionality (*e.g., CD31*, *vWF*, *endothelin-1*, *ICAM2*, and *CXCR4*). Importantly, repressive H3K27me3 marks on N-DVP were increased relative to primed DVP for the non-vascular lineage muscle-specific promoter MYOD1. qRT-PCR expression analysis of these transcripts confirmed that naïve VP indeed expressed significantly higher levels of these vascular genes and lower levels of non-vascular genes (*e.g., PAX6, MSX2, MYOD1*) (Fig. 8e). These results were consistent with an improved epigenetic state in N-DhiPSC that potentiated a lower transcriptional barrier for generating N-DVP with higher vascular-specific gene expressions, decreased non-vascular lineage-primed gene expressions, and ultimately, presumptively greater *in vivo* functionality.

## DISCUSSION

To date, there has not been a human naïve pluripotent stem cell system demonstrating improved effectiveness over conventional hPSC for pre-clinical cellular therapies. These studies describe for the first time the advantage of employing an alternative tankyrase inhibitor-regulated human naïve pluripotent state for improving vascular regenerative therapies. Tankyrase inhibitor-regulated N-hiPSC represent a new class of human stem cells for regenerative medicine with improved multi-lineage functionality. In contrast, conventional hiPSC cultures adopt transcriptomic, epigenetic, and signaling signatures of lineage-primed pluripotency, and display a heterogeneous propensity for lineage bias and differentiation.

Herein, we demonstrated that N-VP differentiated from both normal and diabetic patient-specific N-hiPSC maintained improved genomic stability, possessed higher expressions of vascular identity markers, and decreased expressions of non-vascular lineage-primed genes than VP generated from conventional, primed hiPSC. Moreover, N-DVP were functionally superior in migrating to and re-vascularizing the deep neural layers of the ischemic retina than DVP generated from conventional DhiPSC. Embryonic N-VP with prolific endothelial-pericytic potential and improved vascular functionality for re-vascularizing ischemia-damaged tissues can be generated in unlimited quantities and injected at multiple target sites for multiple treatments and time periods. Such epigenetically plastic N-VP are non-existent in circulating adult peripheral blood or bone marrow. For example, adult EPC are limited in multipotency, expansion, homing, and functionality in diabetes ^2, 14–16^. The generation of embryonic N-DVP from a diabetic patient bypasses this obstacle. N-DhiPSC are more effectively reprogrammed from a donor’s skin or blood cells back to a pre-diseased state, and could subsequently be differentiated to unlimited quantities of pristine, transplantable N-DVP; which unlike adult diabetic EPC would be unaffected by the functional and epigenetic damage caused by chronic hyperglycemia.

Our previous studies demonstrated that embryonic VP derived from conventional CB-derived hiPSC generated with higher and more complete reprogramming efficiencies had decreased lineage-primed gene expression and displayed limited but long-term regeneration of degenerated retinal vessels ^7^. In comparison, conventional skin fibroblast-derived hiPSC lines with higher rates of reprogramming errors and lineage-primed gene expression displayed poorer vascular differentiation and *in vivo* retinal engraftment efficiencies relative to conventional CB-hiPSC. Here, this obstacle was solved for diabetic skin fibroblast donor-derived hiPSC by demonstrating that CD31^+^CD146^+^ endothelial-pericytic N-DVP were more efficiently generated from N-DhiPSC than from conventional DhiPSC. Additionally, N-DVP had higher epigenomic stability, reduced lineage priming, and improved *in vivo* engraftment capacity in ischemia-damaged blood vessels. In future clinical studies, multiple cell types (*e.g*., vascular endothelium, pericytes, retinal neurons, glia, and retinal pigmented epithelium) could all potentially be differentiated from the same autologous or HLA-compatible, banked patient-specific hiPSC line for a comprehensive repair of ischemic vascular and macular degenerative disease.

The studies herein have also demonstrated that the obstacles of incomplete reprogramming, lineage priming, and disease-associated epigenetic aberrations in conventional hiPSC can be overcome with molecular reversion to a tankyrase inhibitor-regulated naïve epiblast-like state with a more primitive, unbiased epigenetic configuration (Fig. 9a). N-DhiPSC possessed a naïve epiblast-like state with decreased epigenetic barriers for vascular lineage specification, and decreased non-vascular lineage specific gene expression. Interestingly, compared to conventional lineage primed DhiPSC, tankyrase-inhibited N-DhiPSC possessed a de-repressed naïve epiblast-like epigenetic configuration at bivalent developmental promoters that was highly poised for non-biased, multi-lineage lineage specification, and was configured in a manner akin to naïve murine ESC ^38–41^ (Fig. 9b).

**Figure 9.**
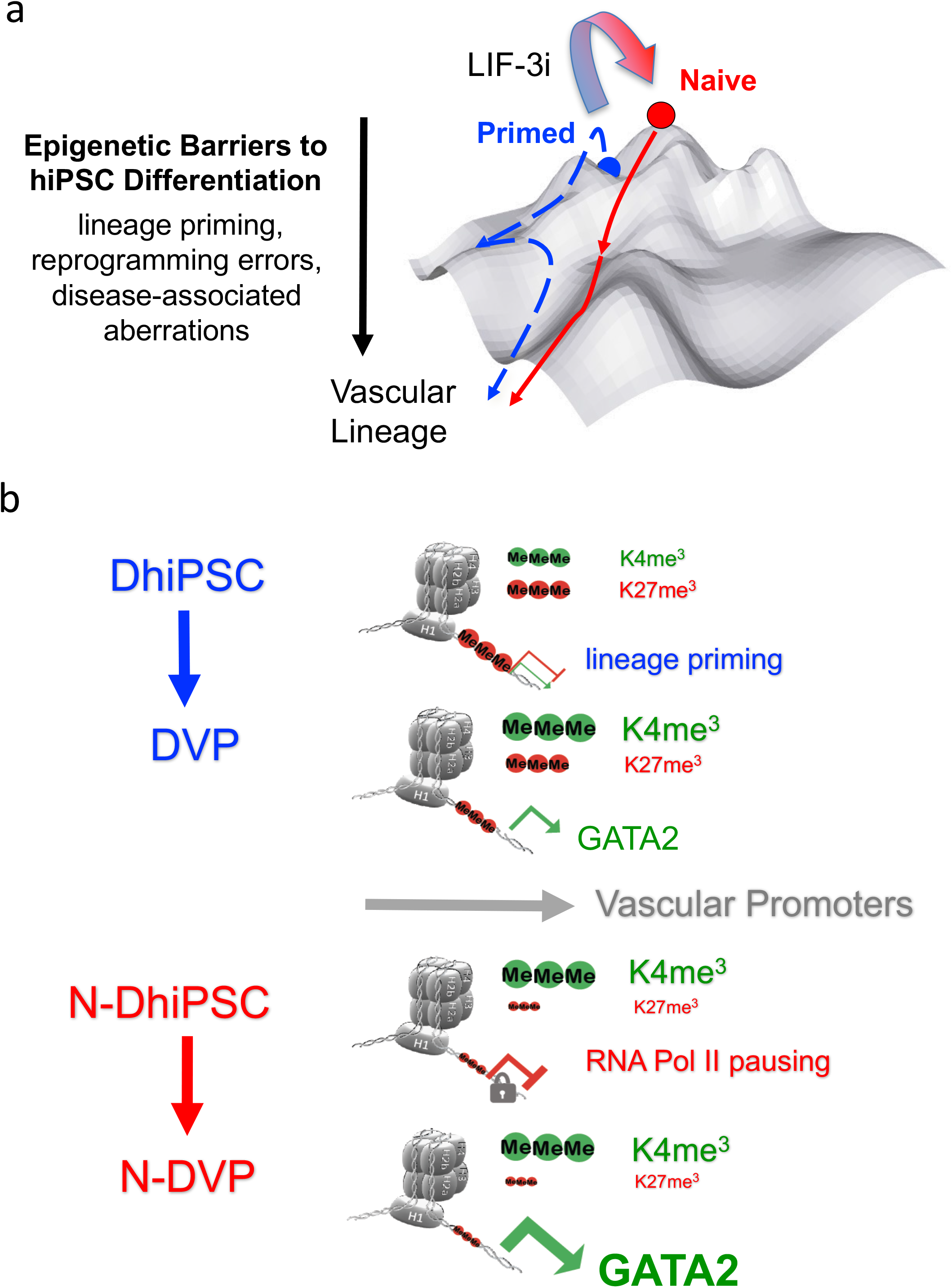
Models for tankyrase inhibitor-mediated reversion of conventional primed hiPSC to an epigenetically plastic naive pluripotent state. (**a**) the epigenetic obstacles of incomplete reprogramming, lineage priming, and disease-associated epigenetic aberrations in conventional primed hiPSC (blue) can be overcome with molecular reversion to a tankyrase inhibitor-regulated naïve epiblast-like state (red) with a more primitive, unbiased epigenetic configuration. (**b**) Compared to conventional lineage primed DhiPSC, tankyrase-inhibited N-DhiPSC possessed a de-repressed naïve epiblast-like epigenetic configuration at bivalent PRC2-regulated developmental promoters that was highly poised for non-biased, multi-lineage lineage specification.

Zimmerlin *et al* first reported that continuous culture of conventional hiPSC with 2i (GSK3β/MEK inhibitors CHIR99021 and PD0325901) plus only the tankyrase/PARP inhibitor XAV939 (LIF-3i) was sufficient for reversion of conventional hPSC to a naïve-like pluripotent state with significantly enhanced developmental potency to all three embryonic germ layer lineages ^12^. Interestingly, several recent studies also incorporated tankyrase/PARP inhibition into their small molecule cocktails to significantly improve the functionality of either murine PSC ^19^, or alternatively human PSC ^21, 45^ with a non-naive epiblast-like epigenetic phenotype. For example, the LCDM ^45^ and mouse expanded potential stem cell (EPSC) ^19, 21^ approaches incorporated either tankyrase or PARP inhibiters in their chemical cocktails to improve both trophectoderm and embryonic contribution of mESC into murine chimeras. Both systems utilized tankyrase/PARP inhibition either at the initiation ^45^ or throughout ^19, 21^ PSC expansion. Remarkably, tankyrase/PARP inhibition of mESC and hPSC preserved the capacity of cleavage-stage murine blastomeres for trophectoderm/ICM lineage segregation. Supplementation with XAV939 to other human naïve-like pluripotent states prior to directed differentiation also led to a significant reduction of lineage-primed gene expressions, and partially rescued a multi-lineage differentiation block ^46^. Unlike the original LIF-3i method, subsequent tankyrase-inhibited human PSC methods did not continuously incorporate a MEK inhibitor in their initial growth conditions, which may be necessary to epigenetically potentiate and maintain a naïve epiblast-like pluripotent state ^38, 40^. LIF-3i was sufficient for bulk, stable reversion and expansion of a large repertoire of conventional lineage-primed hiPSC to a human naïve pluripotent state that possessed characteristics of human preimplantation ICM and mESC, including high clonal proliferation (*i.e*., without requirement of apoptosis inhibition; Rock/Rho kinase inhibitor), MEK-ERK/bFGF signaling independence, activated JAK/STAT3 phosphorylation/signaling, expression of human preimplantation epiblast-like core pluripotency circuits, distal OCT4 enhancer usage, global DNA CpG hypomethylation, and increased expressions of activated β-catenin. Moreover, tankyrase inhibitor-based naïve reversion produced N-hiPSC that retained normal patterns of genomic CpG methylated imprints, reduced lineage-primed gene expression, improved multi-lineage differentiation potency, and did not require reversion culture back to primed conditions prior to differentiation.

The mechanism by which the tankyrase/PARP inhibitor XAV939 stabilized and expanded the functional pluripotency of an inherently unstable human naïve state in classical 2i conditions currently remains incompletely defined. However, herein we suggested a potential epigenetic mechanism that CpG DNA methylation and histone configurations at developmental promoters of diabetic N-hiPSC possessed tight regulation of lineage-specific gene expression and a de-repressed naïve epiblast-like epigenetic state that was highly poised for multi-lineage transcriptional activation. Furthermore, as previously described ^12^, the LIF-3i chemical cocktail minimally employs MEK inhibition (PD0325901) to block lineage-primed differentiation, along with a simultaneous and parallel dual synergy of XAV939 with the GSK3β (CHIR99021) inhibitor to augment WNT signalling ^20^. The presumptive mechanism of augmented WNT signalling is via inhibition of tankyrase-mediated degradation of AXIN, which causes stabilization and increased cytoplasmic retention of the activated isoform of β-catenin in murine ESC (which decreases β-catenin-TCF interactions). However, in humans, the repertoire of proteins directly targeted by tankyrase post-translational PARylation extends far beyond WNT signalling, and includes proteins (*e.g*., AXIN1 and 2, APC2, NKD1, NKD2, and HectD1) with diverse biological functions that potentially cooperate to support a stable pluripotent state ^47^. These functions include regulation of telomere elongation and cohesion (TRF1), YAP signalling (angiomotin), mitotic spindle integrity (NuMa), GLUT4 vesicle trafficking (IRAP), DNA damage response regulation (CHEK2), and microRNA processing (DICER). Interestingly, TRF1 was identified as an essential factor for iPSC reprogramming in mouse and human PSC ^48^. Additionally, although LIF-3i includes MEK inhibition and promotes global and genome-wide low DNA methylation, and, it does not appear to impair genomic CpG methylation at imprinted loci ^12^. Although the mechanism of such imprint preservation by XAV939 in the context of MEK inhibitor is currently obscure, PARylation has been shown to safeguard the Dnmt1 promoter in mouse cells, and antagonizes aberrant hypomethylation at CpG islands, including at imprinted genes ^49, 50^. Thus, the role of PARylation on DNA methylation requires deeper investigation.

Diabetic hyperglycemic alterations of blood vessel viability and integrity lead to multi-organ dysfunction that results in endothelial dysfunction linked to epigenetic remodeling ^17^ (*e.g*., DNA methylation ^51^, histone marks ^52, 53^ and oxidative stress ^54, 55^). Several studies have shown that these aberrant epigenetic changes may be partially overcome by genome-wide chemical treatments that restore some endothelial function ^56, 57^. The extent of retention of diseased ‘diabetic epigenetic memory’ at developmental genes from incomplete or ineffective reprogramming within DhiPSC-derived lineages and its role in impaired regenerative capacity remains unclear, and marked by high variability in differentiation efficiency or retention of diseased phenotype ^58–61^. For example, endothelial differentiation of iPSC generated from diabetic mice displayed vascular dysfunction, impaired *in vivo* regenerative capacity, and diabetic iPSC displayed poor teratoma formation ^63^. Human iPSC from patients with rare forms of diabetes-related metabolic disorders have similarly shown significant functional endothelial impairment ^58^. Transient chemical demethylation of T1D-hiPSC was sufficient to restore differentiation in resistant cell lines and achieve functional differentiation into insulin-producing cells ^18^.

In summary, these studies have demonstrated that highly functional N-VP cells can be generated independent of genetic background or diseased origin from a diseased N-hPSC. Naïve reversion of conventional DhiPSC may potentiate an epigenetic remodeling of reprogrammed diabetic fibroblasts that avoided differentiation into dysregulated in dysfunctional ECs with ‘diabetic epigenetic memory’. Similarly, tankyrase inhibitor-regulated N-DhiPSC are expected to improve the poor and variable DhiPSC differentiation generation of other affected tissues in diabetes ^64^ including pancreatic, renal, hematopoietic, retinal, and cardiac lineages. We propose that autologous or cell-banked transplantable progenitors derived from tankyrase inhibitor-regulated N-hiPSC will more effectively reverse the epigenetic pathology that drive diseases such as diabetes. The application of this new class of human stem cells may inspire further new directions of investigation for understanding human pluripotency, and for improving the utility of hiPSC therapies in regenerative medicine. The further optimization of tankyrase-inhibited human naïve pluripotent stem cells in defined, clinical-grade conditions may significantly advance regenerative medicine.

## METHODS

### Bioethics

hESC lines used in these studies as controls for hiPSC were obtained commercially from the Wisconsin International Stem Cell Bank (WISCB). All hESC experiments proposed conform to guidelines outlined by the National Academy of Sciences, and the International Society of Stem Cell Research (ISSCR). Commercially-acquired hESC are under purview of the Johns Hopkins University (JHU) Institutional Stem Cell Research Oversight (ISCRO), and conform to Institutional standards regarding informed consent and provenance evaluation. All experiments proposed received approval by the JHU ISCRO committee. All animal use and surgical procedures were performed in accordance with protocols approved by the Johns Hopkins School of Medicine Institute of Animal Care and Use Committee (IACUC) and the Association for Research of Vision and Ophthalmology statement for the Use of Animals in Ophthalmic and Visual Research.

### Conventional primed (E8) and naïve (LIF-3i) cultures of hESC and hiPSC

All human embryonic stem cells (hESC) and hiPSC lines used in these studies (**Table S1**) were maintained and expanded in undifferentiated conventional feeder-free primed states in Essential 8 (E8) medium, or naive-reverted with the LIF-3i system, as described ^12, 13^.

Conventional cultures of hiPSC were propagated using commercial E8 medium (ThermoFisher Scientific), or an in-house variant formulation consisting of DMEM/F-12 supplemented with 2.5mM L-Glutamine, 15mM HEPES and 14mM sodium bicarbonate (ThermoFisher Scientific, cat# 11330), 50-100ng/mL recombinant human FGF-basic (Peprotech), 2ng/mL recombinant human TGF-β1 (Peprotech), 64μg/mL L-ascorbic acid-2-phosphate magnesium (Sigma), 14ng/mL sodium selenite (Sigma), 10.7 μg/mL recombinant human transferrin (Sigma), and 20μg/mL recombinant human insulin (Peprotech). Conventional hiPSC were expanded in E8 onto Vitronectin XF (STEMCELL Technologies) matrix-coated tissue culture-treated 6-well plates (Corning). E8 medium was replaced daily and hiPSC were gently passaged every 5-6 days by mechanical selection or bulk passaged using non-enzymatic reagents (*i.e*., Versene solution (ThermoFisher Scientific) or Phosphate-Buffer-Saline (PBS)-based enzyme-free cell dissociation buffer (ThermoFisher Scientific, #13151).

LIF-3i medium was prepared fresh every other week and consists of DMEM/F-12 supplemented with 20% KnockOut Serum Replacement (KOSR, ThermoFisher Scientific), 0.1mM MEM non-essential amino acids (MEM NEAA, ThermoFisher Scientific), 1mM L-Glutamine (ThermoFisher Scientific), 0.1mM β-mercaptoethanol (Sigma), 20ng/mL recombinant human LIF (Peprotech), 3µM CHIR99021 (Tocris or Peprotech), 1µM PD0325901 (Sigma or Peprotech), and 4µM XAV939 (Sigma or Peprotech). Prior to switching between E8 and LIF-3i media, hPSC were adapted for one passage in LIF-5i, as described ^12, 13^. LIF-5i was prepared by supplementing LIF-3i with 10µM Forskolin (Stemgent or Peprotech), 2µM purmorphamine (Stemgent or Peprotech) and 10ng/mL recombinant human FGF-basic (Peprotech). Briefly, primed hiPSC were adapted overnight by substituting E8 with LIF-5i medium. The next day, hiPSC were enzymatically dissociated (Accutase, ThermoFisher Scientific) and transferred onto irradiated mouse embryonic fibroblast (MEF) feeders in LIF-5i medium for only one passage (2 to 3 days). All subsequent passages were grown in LIF-3i medium on MEF feeders. Isogenic E8 cultures were maintained in parallel for simultaneous phenotypic characterization, as previously described in detail ^12, 13^.

### Reprogramming of diabetic fibroblasts to conventional DhiPSC

Adult human Type-I diabetic (T1D) fibroblasts obtained with patients’ informed consent, were purchased from DV Biologics, and cultured in fibroblast culture medium (I-Gro medium, DV Biologics). For reprogramming, single cells were obtained using Accutase and counted. Episomal expression of seven genes (*SOX2*, *OCT4*, *KLF4*, *c-MYC*, *NANOG*, *LIN28*, *SV40LT*) was accomplished by nucleofection of 1×10^6^ diabetic fibroblast cells with 2 μg each of three plasmids, pCEP4-EO2S-EN2L, pCEP4-EO2S-ET2K, and pCEP4-EO2S-EM2K as described^27, 28^. Fibroblasts were nucleofected using human dermal fibroblast nucleofector kits (Lonza, VPD-1001) with Amaxa nucleofector program U-023. Nucleofected cells were transferred onto irradiated MEF in fibroblast growth medium supplemented with 10 μM Rho-associated, coiled-coil containing protein kinase (ROCK) inhibitor Y27362 (Stemgent). The next day, 2 mL of DMEM/F-12 supplemented with 20% KOSR, 0.1mM MEM NEAA, 1mM L-Glutamine, 0.1mM β-mercaptoethanol, 50 ng/mL bFGF, 10 μM Y27362, 5 μg/mL ascorbic acid, and 3 μM CHIR99021 was added. Half of the medium was replaced with fresh medium without Y27362 every other day, until hiPSC colonies appeared. Individual hiPSC colonies were manually isolated, further expanded onto vitronectin-coated plates in E8 medium, or cryopreserved.

### Parallel isogenic primed hiPSC *vs.* N-hiPSC directed neuroectodermal, endodermal, and vascular differentiations *in vitro*

To examine the differentiation competence of normal and diabetic N-hiPSC, we directly differentiated LIF-3i-reverted naïve vs their primed genotypically-identical isogenic (same line) sibling hiPSC counterparts in parallel, as previously described without additional cell culture manipulations ^12, 13^. “Re-priming” (*i.e.,* converting N-hiPSC back to conventional primed conditions prior to their use in directed differentiation assays ^25, 26^) was not necessary with the LIF-3i method ^12, 13^. To minimize hiPSC assay variations within directed differentiation experiments that may arise from hiPSC interline variability and genetic background bias, paired isogenic primed and LIF-3i-reverted hiPSC lines were simultaneously and directly cultured into defined, identical, feeder-free differentiation systems according to manufacturer’s directions. Naïve reversions were performed in LIF-5i/LIF-3i media fresh for each differentiation experiment starting from a low passage primed hPSC line, as described ^13^. For example, for functional comparisons of naïve vs. primed isogenic hiPSC lines, sibling cultures were prepared at equivalent passage number, starting from the primed parental hPSC line. Primed and naïve hPSC sibling cultures were expanded in parallel in their respective media for 5-7 passages before differentiation (*e.g*., E8 *vs*. LIF-3i, see schematic Fig 3a). This experimental approach for primed vs naïve differentiation was previously employed for functional comparison of primed vs naïve hiPSC states ^12, 13^. Detailed information for the origins and derivation of all hiPSC lines used in these studies for these assays was previously reported ^12^ **(Table S1)**.

Differentiation to neural progenitors was performed using GIBCO PSC neural induction medium (NIM; ThermoFisher Scientific, A1647801) and the manufacturer’s recommendations. Differentiation into definitive endodermal progenitors was achieved using the StemDiff Definitive Endoderm Kit (StemCell Technologies) following manufacturer’s protocols. Vascular differentiation of VP from primed and naïve hiPSC was modified and optimized from methods we previously described ^7^. The experimental approach is summarized in FIG. 3a. Briefly, VP differentiation was performed using a modified protocol based on the STEMdiff APEL-Li medium system ^32^. Briefly, APEL-2Li medium (StemCell Technologies, #5271) was supplemented with Activin A (25 ng/mL), VEGF (50 ng/mL), BMP4 (30 ng/mL), and CHIR99021 (1.5 μM) for the first 2 days, and then APEL-Li that was supplemented with VEGF (50 ng/mL) and SB431542 (10μM). Differentiation medium was replaced every 2 days until cells were harvested for analysis.

### Isogenic primed *vs.* naive hiPSC teratoma assays

Isogenic (genotypic-identical; same line) primed (E8) and naïve (LIF-3i) hiPSC cultures were maintained in parallel for 9 passages prior to teratoma formation assays. Teratomas were directly generated from a fixed number of cells (5×10^6^) and duration (8 weeks) in isogenic primed *vs*. LIF-3i naïve hiPSC conditions. LIF-3i-cultured N-hiPSC colonies did not require chemical manipulation or re-priming culture steps prior to enzymatic harvest from culture and direct injection into NOG mice. Adherent primed vs naive hiPSC were collected using Accutase and counted using Countess counter (ThermoFisher Scientific). For all experiments, 5×10^6^ hiPSC were admixed with Growth factor reduced Matrigel (Corning, cat# 356230) on ice. Cells were injected subcutaneously into the hind limbs of immunodeficient NOG male sibling mice. Teratomas were dissected 8 weeks following injection and fixed by overnight immersion in PBS, 4% formaldehyde. All tissues were paraffin-embedded, and microsectioned (5μm thickness) onto microscope glass slides (Cardinal Health) by the Histology laboratory from the Pathology Department at the Johns Hopkins University. To account for heterogeneous teratoma histological distribution, 15 individual equally-spaced sections were immunostained per tissue for each antigen of interest and quantification. Slides were heated in a hybridization oven (ThermoFisher Scientific) at 60°C for 20 minutes and then kept at room temperature for 1 hour to dry. Paraffin was eliminated by three consecutive immersions in xylenes (Sigma) and sections were rehydrated by transitioning the slides in successive 100%, 95%, 70% and 0% ethanol baths. Sections were placed in 1X wash buffer (Dako) prior to heat-induced antigen retrieval using 1X Tris-EDTA, pH9 target retrieval solution (Dako) and wet autoclave (125°C, 20 min). Slides were cooled and progressively transitioned to PBS. After 2 washes, tissues were blocked for 1 hour at room temperature using PBS, 5% goat serum (Sigma), 0.05% Tween 20. Endogenous biotin receptors and streptavidin binding sites were saturated using the Streptavidin/Biotin Blocking kit (Vector Laboratories). All antibodies were diluted in blocking solution. Sections were incubated overnight at 4°C with monoclonal mouse anti-NG2 (Sigma, C8035, 1:100), mouse anti-SOX2 (ThermoFisher Scientific, MAS-15734, 1:100) or rabbit anti-cytokeratin 8 (Abcam, ab53280, 1:400) primary antibodies, washed 3 times, incubated for 1 hour at room temperature with biotinylated goat anti-mouse or goat anti-rabbit IgG antibodies (Dako, 1:500), washed 3 times and incubated with streptavidin Cy3 (Sigma, 1:500) for 30 minutes at room temperature. After 2 washes, tissues were incubated for 2 hours at room temperature with a second primary antibody (*e.g*., anti-Ki67) differing in species from the first primary antibody. After incubation with rabbit (Abcam, ab16667, 1:50) or mouse (Dako, M7240, 1:50) anti-Ki67 monoclonal antibody, sections were washed 3 times and incubated for 1 hour at room temperature with highly cross-adsorbed Alexa Fluor 488-conjugated goat anti-rabbit or goat anti-mouse secondary antibody (ThermoFisher Scientific, 1:250). Sections were washed twice, incubated with 10µg/mL DAPI (ThermoFisher Scientific, D1306) in PBS, washed 3 times in PBS and slides were mounted with coverslips using Prolong Gold Anti-fade reagent (ThermoFisher Scientific) for imaging. Isotype controls for mouse (ThermoFisher Scientific) and rabbit (Dako) antibodies were substituted at matching concentration with primary antibodies as negative controls.

For teratoma organoid quantifications, photomicrographs were obtained using a 20X objective and Zeiss LSM 510 Meta Confocal Microscope. Image processing and quantification was performed using NIS-Elements software (Nikon). The ROI editor component was applied to autodetect regions of interest in the Cy3 channel that delineated lineage-defined structures (i.e., Cytokeratin 8^+^ definitive endoderm, NG2^+^ chondroblasts, SOX2^+^ neural rosettes) within teratomas. Thresholding and restrictions were standardized in the Object Count component and applied to detect and export the number of DAPI^+^ and Ki67^+^ nuclei within ROIs for all analyzed sections.

### Antibodies

Source and working dilutions of all antibodies used in these studies for Western blots, FACS, genomic dot blots, ChIP, and immunofluorescence experiments are listed in **Table S3.**

### Western blotting

Cells were collected from either primed (E8 media on vitronectin-coated plates) or naïve (LIF-3i/MEF plates) conditions with Enzyme-Free Cell Dissociation Buffer (Gibco, 13151-014). Cells were washed in PBS and pelleted. Cell pellets were lysed in 1x RIPA buffer (ThermoFisher Scientific, 89900), 1mM EDTA, 1x Protease Inhibitor (ThermoFisher Scientific, 78430), and quantified using the Pierce bicinchoninic acid (BCA) assay method (ThermoFisher Scientific). 25μg of protein per sample was loaded on a 4-12% NuPage Gel (ThermoFisher Scientific, NP0336) according to manufacturer’s recommendations. The gel was transferred using the iBlot2 (Life Technologies), blocked in Tris-buffered saline (TBS), 5% non-fat dry milk (Labscientific), 0.1% Tween-20 (TBS-T) for 1 hour, and incubated overnight at 4°C in with anti-phosphorylated-STAT3 primary antibody (Cell Signaling, 9145) according to manufacturer’s protocols. Membranes were rinsed 3 times in TBS-T, incubated with horseradish peroxidase (HRP) –linked goat anti-rabbit secondary antibody (Cell Signaling, 7074) for 1 hour at room temperature, rinsed 3 times, and developed using Pierce ECL Substrate (ThermoFisher Scientific, 32106). Chemiluminescence detection was imaged using an Amersham Imager 600 (Amersham). Anti-actin antibody staining was performed for each membrane as a loading control.

### Flow cytometry analysis of vascular differentiations and FACS purification of CD31^+^CD146^+^ endothelial-pericytic VP populations

Recipes for all differentiation reagents, antibodies, and PCR primers were previously described and also summarized in **Table S3**. For flow cytometry analysis of vascular differentiations, cells were washed once in PBS, and enzymatically digested with 0.05% trypsin-EDTA (5 min, 37°C), neutralized with FCS, and cell suspensions were filtered through a 40 μm cell-strainer (Fisher Scientific, Pittsburgh, PA). Cells were centrifuged (200 g, 5 min, room temperature) and re-suspended in staining buffer (EBM alone or 1:1 EMG2:PBS). Single cell suspensions (<1×10^6^ cells in 100 µL per tube) were incubated for 20 min on ice with directly conjugated mouse monoclonal anti-human antibodies and isotype controls. Cells were washed with 3 mL of PBS, centrifuged (300g, 5 min, room temperature), and resuspended in 300 µL of staining buffer prior to acquisition. Viable cells were analyzed (10,000 events acquired for each sample) using the BD CellQuest Pro analytical software and FACSCalibur™ flow cytometer (BD Biosciences). All data files were analyzed using Flowjo analysis software (Tree Star Inc., Ashland, OR).

FACS of primed *vs.* naive VP populations was performed at the Johns Hopkins FACS Core Facility with a FACS Aria III instrument (BD Biosciences, San Jose, CA). Cell suspensions from APEL vascular differentiations were incubated with mouse anti-human CD31-APC (eBioscience, San Diego, CA) and CD146-PE (BD Biosciences) antibodies for 30 min on ice, and FACS-purified for high CD31 and CD146 expression, plated onto fibronectin-coated plates in EGM2, and expanded to 80-90% confluency for 7-9 days prior to in vitro analyses or *in vivo* injections into the eyes of I/R-treated NOG mice.

### Vascular functional assays

The methods for endothelial Dil-acetylated-LDL uptake assays, Matrigel tube quantitation assays, EdU proliferation assays, β-galactosidase senescent assays were all described previously ^7^, and are summarized below briefly. For Dil-Acetylated-Low Density Lipoprotein (Dil-Ac-LDL) uptake assays, FACS-purified CD31^+^CD146^+^ primed vs naïve VP populations were expanded in EGM2 medium ∼7 days to 60 to 70% confluency on fibronectin pre-coated 6-well plates (1-1.5 x 10^5^ cells/well) prior to Dil-Ac-LDL uptake assays (Life Technologies, Cat No. L-3484). Fresh EGM2 medium supplemented with 10 μg/mL Dil-Ac-LDL, was switched before assays, and incubated for 4 hours at 37°C. Cells were washed in PBS and Dil-Ac-LDL-positive cells imaged with a Nikon Eclipse Ti-u inverted microscope (Nikon Instruments Inc., Melville, NY) and Eclipse imaging software. Cells were also harvested with Accutase (5 min, 37°C) and Dil-Ac-LDL^+^ cells quantitated by flow cytometry.

*In vitro* vascular functionality of primed DVP vs N-DVP was determined with quantitative Matrigel vascular tube-forming assays as previously described **7**. Briefly, MACS-purified CD31^+^CD146^+^ isogenic DVP were expanded in EGM2 on fibronectin-coated (10 μg/mL). tissue culture plates. Adherent cells were treated with Accutase for 5 min, and collected into single cell suspensions. Primed DVP or N-DVP cells were transferred in 48-well plates (2×10^5^ cells/ well in EGM2 medium) pre-coated with Matrigel (Corning, #356237, 200 μL/well). The next day, multiple phase contrast pictures of vascular tube formations were imaged with an inverted Eclipse Ti-u Nikon microscope (Nikon Instruments Inc., Melville, NY) and Eclipse imaging software without overlapping the imaged regions. All the vascular tubes formed by VP, DVP, and N-DVP were measured by Eclipse imaging software. Statistical comparisons were performed with unpaired t-tests using Prism (GraphPad Software, San Diego, CA).

For senescence assays, naïve vs primed VP populations were plated onto fibronectin (10 μg/mL)-coated 6-well tissue culture plates, and VP were expanded in EGM2 for up to 30 days (3-6 passages), and senescent cells were assayed for acidic senescence-associated ß-galactosidase activity. Cells were grown to ∼60-80% confluency in 12-well fibronectin-coated plates prior to analysis. Cells were fixed in 2% paraformaldehyde and β-galactosidase activity was quantified by detecting hydrolysis of the X-gal substrate by colorimetric assay as per manufacturer’s protocol for detection of senescent cells. (Cell Signaling Technology, Danvers, MA). Nuclei were counterstained using the fluorescent dye Hoechst 33342 (BD Biosciences). Total number of Hoechst^+^ cells and blue X-Gal^+^ senescent cells were automatically enumerated using an inverted Eclipse Ti-u Nikon microscope and the Object Count component of the NIS Elements software. For each sample, 2 individual wells were photographed at five independent locations using a 20x objective.

### Transmission Electron Microscopy (TEM) of DVP and N-DVP

Primed vs naïve VP were plated onto fibronectin (10 μg/mL) coated Labtek chambers, culture expanded in EGM2, and fixed for TEM as previously described at the Wilmer Microscopy Core **7**. Sections were imaged with a Hitachi H7600 TEM at 80KV (Gaithersburg, MD) and a side mount AMT CCD camera (Woburn, Mass).

### NCS DNA damage response assays

Primed DhiPSC and N-DhiPSC isogenic (same lines at same passage) were simultaneously differentiated in parallel into DVP and N-DVP using APEL medium, as described above. CD31^+^CD146^+^ VP cells were expanded in EGM2 (3 passages) onto fibronectin-coated (10μg/mL) 6-well plates (for Western blot analysis), or alternatively the last passage was transferred onto 8-well Nunc Labtek II chamber slides for immunostaining. To induce DNA damage, expanded DVP and N-DVP cells were incubated for 5 hours in EGM2 supplemented with 100 ng/mL of the radiomimetic agent neocazinostatin (NCS, Sigma). Untreated DVP and N-DVP cells were analyzed in parallel as controls. Western blot analysis was performed as described above. For detection of phosphorylated H2AX by immunofluorescence, VP cells were fixed for 10 minutes using 1% paraformaldehyde in PBS. For immunofluorescent staining of chambered slides, fixed cells were blocked for non-specific staining and permeabilized using a blocking solution consisting of PBS, 5% goat serum (Sigma) and 0.05% Tween 20 (Sigma). Samples were incubated overnight at 4°C with a rabbit anti-human phospho-H2AX antibody (Cell Signaling, #9718) diluted (1:200) in blocking solution. The next day, VP cells were washed (Dako wash buffer, Dako) and incubated for 2 hours at room temperature with a biotinylated goat anti-rabbit secondary antibody (Dako, 1:500 in blocking solution). Cells were washed 3 times and incubated for 30 minutes with streptavidin Cy3 (Sigma, 1:500). All samples were sequentially washed and incubated with a mouse monoclonal anti-human CD31 (Dako, M0823, 1:100) and Alexa488-conjugated goat anti-mouse secondary antibody (ThermoFisher, 1:100), both for 1 hour at room temperature. Finally, slides were washed in PBS and incubated with DAPI (1:2000) for 5 minutes at room temperature for nuclear staining. Slides were mounted using the Prolong Gold anti-fade mounting reagent (ThermoFisher) and cured overnight. For each condition, 5 to 6 independent frames were captured for the Cy3, Alexa488 and DAPI channels using a 20X objective and a LSM510 Meta confocal microscope (Carl Zeiss Inc., Thornwood, NY) in the Wilmer Eye Institute Imaging Core Facility. Quantification of phospho-H2AX+ foci within DAPI^+^ nuclei of CD31^+^ VP was performed using the NIS-Elements software. Briefly, thresholds and masks were sequentially created for the Alexa488 (CD31) and DAPI channels to limit the analysis to nuclei of VP cells. Nuclei were further defined using the size/area and circularity parameters. Each individual CD31^+^ nucleus was characterized as a single object using the “object count” function. Finally, the number of foci per nucleus was determined by counting the number of objects in the Cy3 channel. A total of 128 to 165 nuclei were analyzed for each condition (primed *vs.* naive ± NCS).

### Ocular I/R Injury and VP Injections into NOD/Shi-*scid*/IL-2Rγ^null^ (NOG) eyes

The I/R ocular injury model was previously described ^7^. Briefly, six-to eight-week old male NOG mice (Johns Hopkins Cancer Center Animal Facility) were subjected to high intraocular pressure to induce retinal ischemia-reperfusion injury. Mice were deep anesthetized by intraperitoneal (IP) injection of ketamine/xylazine (50 mg/kg ketamine + 10 mg/kg xylazine in 0.9% NaCl). The pupils were dilated with 2.5% phenylephrine hydrochloride ophthalmic solution (AK-DILATE, Akorn, Buffalo Glove, IL) followed by 0.5% tetracaine hydrochloride ophthalmic topical anesthetic solution (Phoenix Pharmaceutical, St. Joseph, MO). The anterior chamber of the eye was cannulated under microscopic guidance (OPMI VISU 200 surgical microscope, Zeiss, Gottingen, Germany) with a 30-gauge needle connected to a silicone infusion line providing balanced salt solution (Alcon Laboratories, Fort Worth, TX); avoiding injury to the corneal endothelium, iris, and lens. Retinal ischemia was induced by raising intraocular pressure of cannulated eyes to 120 mmHg for 90 min by elevating the saline reservoir. Ischemia was confirmed by iris whitening and loss of retinal red reflex. Anesthesia was maintained with two doses of 50 µL intramuscular ketamine (20 mg/mL) for up to 90 min. The needle was subsequently withdrawn, intraocular pressure normalized, and reperfusion of the retinal vessels confirmed by reappearance of the red reflex. The contralateral eye of each animal served as a non-ischemic control. Antibiotic ointment (Bacitracin zinc and Polymyxin B sulfate, AK-Poly-Bac, Akron) was applied topically. Two days later, MACS-purified and expanded human DVP and N-DVP were injected into the vitreous body (50,000 cells in 2 µL/eye), using a micro-injector (PLI-100, Harvard Apparatus, Holliston, MA).

### Immunofluorescence staining of flat whole-mounted NOG mouse retinae

Human cell engraftment into NOG mouse retinae was detected directly with anti-human nuclear antigen (HNA) immunohistochemistry with murine vascular marker co-localization (murine CD31 and collagen IV) using anti-murine CD31 and anti-murine collagen IV antibodies. Animals were euthanized for retinal harvests and HNA-positive cell quantitation at 1, 3, and 4 weeks following human VP injection (2 days post-I/R injury). After euthanasia, eyes were enucleated, cornea and lens were removed, and the retina was carefully separated from the choroid and sclera. Retinae were fixed in 2% paraformaldehyde in TBS for overnight at 4°C, and permeabilized via incubation with 0.1% Triton-X-100 in TBS solution for 15 min at 4°C. Following thorough TBS washes, free floating retinas were blocked with 2% normal goat serum in TBS with 1% bovine serum albumin and incubated overnight at 4°C in primary antibody solutions: rabbit anti-mouse Collagen IV (AB756P, Millipore, 1:100) and/or rat anti-mouse CD31 (550274, BioSciences, 1:50) in 0.1% Triton-X-100 in TBS solution (to label basement membrane and EC of blood vessels, respectively). On the next day, retinae were washed with TBS, and incubated with secondary antibodies for 6 hours at 4°C. A goat anti-rabbit Cy3-conjugated secondary antibody (Jackson Immuno Research, # 111-165-003, 1:200) was used to detect collagen IV primary antibody, and a goat anti-rat Alexafluor-647-conjugated secondary antibody (Invitrogen, # A21247, 1:200) was used to detect the anti-CD31 primary antibody. Human cells were detected using directly Cy3-conjugated anti-HNA (Millipore, MAB1281C3, 1:100). After washing in TBS, flat mount retinas were imaged with confocal microscopy (LSM510 Meta, Carl Zeiss Inc., Thornwood, NY) at the Wilmer Eye Institute Imaging Core Facility.

### Immunofluorescent confocal microscopy and quantitation of human cell vascular engraftment in murine retinae

For quantification of HNA^+^ cells in the superficial layers of whole retinae, whole mount retinas were prepared from the eyes of animals at 1, 3 or 4 weeks following intra-vitreal transplantation of human cells (50,000 primed DVP or N-DVP cells per eye) following I/R injury. Non-I/R injured eyes and control PBS-injected eyes were also analyzed as controls. Images were acquired with ZEN software using a 10X objective and a LSM510 Meta confocal microscope. For each individual eye, the entire retina was tile-scanned and stitched (7×7 frames, 10% overlapping).

For human HNA^+^ cell quantification analysis, photomicrographs were processed using the Fiji distribution of imageJ. Briefly, a region of interest was created using the DAPI channel and the “magic wand” function to conservatively delineate the whole retina and exclude from the analysis the limited background at the edges of the retina preparation that could be detected in the Cy3 (HNA) channel for some samples. The Cy3 channel was processed with the “smooth” function and a mask was created using by thresholding. The Cy3 channel was further prepared for the “analyze particle” plugin by using standardized sequential corrections that were limited to despeckle, filtering (Minimum) and watersheding. Particle objects corresponding to HNA+ nuclei were automatically counted using fixed size and circularity parameters.

Eyes were also analyzed for quantification of human CD34^+^ or human CD31^+^ blood vessels within defined layers of the mouse retina in some experiments. Briefly, the anterior eye (cornea/iris) was dissected free by a circumferential cut at the limbus. Eyecups were fixed using paraformaldehyde and prepared for cryopreservation by immersion in gradients of sucrose (Lutty et al IOVS 1993, PMID: 7680639). Eyes were hemisected through the optic nerve (**Fig. S7a**) and the two halves embedded in OCT-sucrose. Serial cryosections (8μm thickness) were prepared from hemisections that included the retina (**Fig S7a**), and stored at −80°C. Equally interspaced microsections [n=11 (E8) and 13 (LIF-3i) for CD34, and n=3 (E8) and 7 (LIF-3i) for CD31 immunostainings]. Retinal sections were sequentially immunostained with either mouse anti-human CD34 (BD Biosciences, clone My10, #347660, 1:50) or mouse anti-human CD31 (Dako, M0823, 1:50) overnight at 4°C followed by goat anti-mouse Alexa488 (ThermoFisher Scientific, 1:200) for 1 hour at room temperature. Sections were subsequently stained with either rabbit anti-mouse collagen type IV (Millipore, AB756P, 1:200) or rat anti-mouse CD31 (BD Biosciences, 550274, 1:50) for 1 hour at room temperature. Alexa647-conjugated goat-anti rabbit (ThermoFisher Scientific, A-21246; 1:200) or goat anti-rat (ThermoFisher Scientific) secondary antibodies (*i.e*., conjugated F(ab’)2 fragments) that were highly cross-adsorbed against IgG from other species were subsequently incubated for 1 hour at room temperature. Nuclei were counterstained using DAPI. Negative immunostaining controls in each experiment were conducted and confirmed negative, and consisted of replacing primary antibodies with mouse, rat and rabbit nonimmune IgG (Dako or ThermoFisher Scientific) at the corresponding antibody concentration to verify absence of unspecific antibody binding. Retinal sections were mounted with Prolong Gold anti-fade reagent (ThermoFisher Scientific) and cured overnight in the dark. Images were acquired using a 20X objective with the ZEN software and a LSM510 Meta confocal microscope.

Photomicrographs were further processed for human cell quantification using the Fiji distribution of imageJ (**Fig. S7b**). Briefly, regions of interest (ROI) were created using the DAPI channel as a template to delineate the GCL, INL, and ONL, the other regions (ILM, IPL, OPL and S) being defined as intercalated around and between the 3 DAPI-defined ROI. Analysis was pursued by processing the Alexa488 (human CD34 or CD31) channel using a sequential series of defined parameters using the “smooth”, “despeckle”, “filter (median)” functions and thresholding. Alexa488^+^ objects were counted within and between ROI using the “analyzes particles” plugin. Human CD34 (**Fig S7b**) or human CD31 expression was scored only when the Alexa488^+^ signal was expressed at chimeric human-murine blood vessels that also expressed murine collagen IV (mColIV) or murine CD31 (mCD31) (**Fig, 7a,c**). The quantity of human blood vessels detected in murine vessels ranged between 1-13 per image at 20X objectives (**Fig, 7b,d**).

### Quantitative Real-time Polymerase Chain Reaction (qRT-PCR) and Chromatin Immunoprecipitation PCR (ChIP-PCR)

The sequences and published reference citations of all PCR primers used in these studies for qRT-PCR and qChIP-PCR are listed in **Table S3.** For qRT-PCR analyses, feeder-dependent LIF-3i hPSC cultures were MEF-depleted by pre-plating onto 0.1% gelatin-coated plates for 1 hour at 37°C, as previously described ^12^. Samples were sequentially and simultaneously collected from representative hPSC lines in primed (E8), or naïve (LIF-3i; p>3) conditions. Alternatively, genotypic-identical (isogenic) paired samples were prepared from EGM2-expanded primed and naïve VP. Total RNA was isolated from snap-frozen samples using the RNeasy Mini Kit (Qiagen) following the manufacturer’s instructions, and quantified using a Nanodrop spectrophotometer (ThermoFisher Scientific). Genomic DNA was eliminated by in-column DNase (Qiagen) digestion. Reverse transcription of RNA (1µg/sample) was accomplished using the SuperScript VILO cDNA Synthesis Kit (ThermoFisher Scientific) and a MasterCycler EPgradient (Eppendorf). For real-time PCR amplification, diluted (1:20) cDNA samples were admixed to the TaqMan Fast Advanced Master Mix (ThermoFisher Scientific) and Taqman gene expression assays (ThermoFisher Scientific).

Matching isogenic samples were prepared in parallel for ChiP-PCR assays. Isogenic hPSC cultures were expanded using primed (E8) and naïve (LIF-3i/MEF) conditions and analyzed at passages matching RT-PCR analysis. Alternatvely, VP cells were prepared from isogenic primed and naïve PSC using the same APEL/EGM2 conditions as the samples prepared for RT-PCR. Cells were collected using Accutase and counted using a Countess cell counter (ThermoFisher Scientific). Feeders were excluded from LIF-3i/MEF samples by pre-plating for 1 hour on gelatin-coated plates and pre-plated samples were re-counted after the pre-plating step. 3 x10^6^ cells were allocated per ChIP assay and prepared using the Magna ChIP A/G chromatin immunoprecipitation kit (Millipore). Cells were centrifuged (300g), supernatant was discarded and cells were fixed for 10 minutes at room temperature by resuspending in 1mL of PBS, 1% formaldehyde (Affymetrix). Unreacted formaldehyde was quenched using 100μL 10X Glycine (Millipore). Samples were left at room temperature for 5 minutes, centrifuged (300g) and washed twice in 1mL ice-cold PBS. Samples were resuspended in ice-cold PBS containing either 1X Protease Inhibitor Cocktail II (Millipore) or 1X complete Mini protease inhibitor (Roche). Samples were centrifuged at 800g for 5 minutes, cell pellets were snap-frozen in liquid nitrogen and stored at −80°C until use for ChIP assay. Cell lysis, homogenization and nuclear extraction of cryopreserved samples were processed using the reagents provided in the Magna ChIP kit and the manufacturer instructions. The isolated chromatin was fragmented using a Diagenode Bioruptor Plus sonication device. Sonication settings (10 cycles, high, 30s on, 30s off) were validated in pilot experiments to shear cross-linked DNA to 200-1000 base pairs by agarose gel electrophoresis. The sheared chromatin was centrifuged at 10,000g at 4°C for 10 minutes and immediately processed for immunoprecipitation. 1×10^6^ cell equivalent of cross-linked sheared chromatin were prepared according to the kit manufacturer’s protocol. Briefly, 1% of sheared chromatin was separated as “input” control. The remaining sample was admixed with 5 μg of immuno-precipitating antibody (**Table S3**) and protein A/G magnetic beads. Antibodies were substituted with corresponding rabbit or mouse IgG (**Table S3**) as negative isotype controls using 5% sheared chromatin. The chromatin-antibody-beads mixture was left incubating overnight at 4°C with agitation. Protein A/G beads were pelleted using a MagJET separation rack (ThermoFisher Scientific) and supernatant was discarded. Protein/DNA complexes were washed and eluted, beads were separated using the MagJET rack and DNA was purified according to the manufacturer’s instructions. The immunoprecipitated genomic DNA was amplified using the Power SYBR Green Master Mix (ThermoFisher Scientific) with relevant published primers for GAPDH, GATA2, GATA6, HAND1, NANOG, MSX2, PAX6, SOX1 (**Table S3**), CD31, vWF, endothelin-1, ICAM2, MYOD1 (all from ^1^), CXCR4, DLL1, FZD7 and ELP3 (all from reference ^65^) (**Table S3**) using a ViAA7 Real Time PCR System (ThermoFisher Scientific). Specificity of antibodies was validated using the isotype controls and samples were normalized to their corresponding input controls.

### Genomic DNA dot-blots of 5-methylcytosine (5MC) and 5-hydroxymethylcytosine (5hMC) CpG methylation

Genomic DNA from isogenic parallel primed (E8) and preplated LIF-3i cultures of representative hiPSC lines was extracted using the DNeasy Blood and tissue Kit (Qiagen) and quantified using a Nanodrop spectrophotometer (ThermoFisher Scientific), as described ^12^. For each sample, 1.6μg DNA was diluted in 50μL of nuclease-free water (Ambion), denatured by adding 50μL of 0.2M NaOH, 20mM EDTA and incubating for 10 minutes at 95°C, and neutralized by adding 100μL 20X Saline-Sodium Citrate SSC hybridization buffer (G Biosciences) and chilling on ice. A series of five 2-fold dilutions (800ng to 50ng) and nuclease-free water controls were spotted on a pre-wetted (10X SSC buffer) nylon membrane using a Bio-Dot Microfiltration Apparatus (Bio-Rad). The blotted membrane was air-dried and UV-cross-linked at 1200 J/m^2^ using a UV Stratalinker 1800 (Stratagene). The membrane was blocked in TBST, 5% nonfat dry milk for 1 hour at room temperature with gentle agitation, washed 3 times in TBST, and incubated at 4°C overnight with rabbit anti-5mC (Cell Signaling, 1:1000) or anti-5hmC (Active Motif, 1:5000) antibodies diluted in TBST, 5% BSA. The membrane was washed 3 times in TBST and incubated for 1 hour at room temperature with HRP-conjugated anti-rabbit secondary antibody (Cell Signaling) diluted 1:1000 in blocking buffer. After 3 washes in TBST, the membrane was treated with Pierce ECL Substrate (ThermoFisher) for chemiluminescent detection with an Amersham Imager 600 (Amersham). After acquisition, the membrane was washed 3 times in H_2_O and immersed in 0.1% methylene blue (Sigma), 0.1M sodium acetate stain solution for 10 minutes at room temperature. Excess methylene blue was washed 3 times in water with gentle agitation. Colorimetric detection was achieved using the Amersham Imager 600 (Amersham). 5mC, 5hmC, and methylene blue intensities were quantified by ImageJ software.

### Statistics

Statistical significance was determined using statistical graphing software (Prism GraphPad) using two-tailed *t* tests (between individual groups), or 1-way analysis of variance (*e.g.,* analysis of variance-Eisenhart method with Bonferroni correction) for statistical testing of ≥3 groups. For smaller, non-Gaussian–distributed sample sizes (n<10), nonparametric (Mann-Whitney) tests were performed. *P* values of at least <0.05 were considered significant.

### Bioinformatics Analyses

The (Illumina, San Diego, CA gene expression arrays (Illumina Human HT-12 Expression BeadChip) and Infinium 450K CpG methylation raw array data analyzed in these studies were published previously and available at Gene Expression Omnibus under accession numbers GSE65211 and GSE65214, respectively, and processed as previously described ^12^. Gene specific enrichment analysis (GSEA) of expression arrays was conducted as described ^66^. The bioinformatics method for calculating crossplots of differential promoter CpG methylation beta values *vs.* corresponding differential gene expression was previously described ^12^.

## ACKNOWLEDGEMENTS

This work was supported by grants from the NIH/NEI (R01EY023962), NIH/NICHD (R01HD082098), Novo-Nordisk Diabetes & Obesity Science Forum Award, RPB Stein Innovation Award, The Maryland Stem Cell Research Fund (2018-MSCRFV-4048, 2014-MSCRFE-118153), an RPB Unrestricted grant (Wilmer), The Lisa Dean Moseley Foundation, and Wilmer core grant for vision research (EY001765). We are grateful for technical support by Schuyler Metzger and Jessica Davidson.

## AUTHOR CONTRIBUTIONS

TSP, LZ, and ETZ designed all experiments and wrote manuscript. TSP, LZ, REM, JT, JSH, RK, AH, NR, SM, RG, and IB performed and analyzed experiments. All authors edited the manuscript, interpreted results, and gave final approval of the manuscript. ETZ supervised and directed the studies.

## COMPETING INTERESTS

Authors do not declare any competing financial interests.

## SUPPLEMENTAL INFORMATION

Accompanies this manuscript at the following weblink:

