## Supplemental Data for "Vascular Progenitors Generated from Tankyrase Inhibitor-Regulated Naïve Diabetic Human iPSC Potentiate Efficient Revascularization of Ischemic Retina"

### A. <u>Supplementary Tables</u>

**Supplementary Table 1 (.xlsx file). Human iPSC lines and naïve reversion methods**

**Supplementary Table 2 (.xlsx file). Human iPSC cell line karyotype summary**

**Supplementary Table 3 (.xlsx file). Antibodies, PCR primers, and lineage-specific genes (PRC2 module)**

### B. Supplementary Figures

**Supplementary Figure 1.** ***In vitro*** **multi-lineage directed differentiations of paired isogenic primed *vs.* naïve normal (non-diabetic) hiPSC lines.** Isogenic primed (E8 cultures) vs. naïve-reverted (LIF-3i cultures) hiPSC were differentiated in parallel using established multi-lineage protocols and commercially available kits, as previously described and without any requirement for re-priming [12,13]. Shown are direct comparisons between normal (non-diabetic) isogenic hiPSC lines demonstrating augmented differentiation capacities to all three germ layers and robust/improved capacity for terminal differentiation. (**a)** Neuro-ectodermal differentiation of isogenic comparisons of primed *vs*. naïve hPSC lines performed as described with PSC neural induction medium [12] (Thermo Fisher Scientific; *n*=6, three normal fibroblast-hiPSC lines (circles; C1.2, C2, 7ta) and three normal cord blood (CB)-derived hiPSC lines (triangles; E5C3, E5C1, LZ610); see **Table S1**). Shown are flow cytometry analysis of differentiation cultures for protein expressions of % Nestin$^+$SOX1$^+$ neural progenitor cells determined by flow cytometry as described [12,13]. (**b**) Representative images of Tuj1-expressing terminally-differentiated cells from a primed (E8 culture) vs naïve (LIF-3i culture) CB-iPSC line (E5C3) after 5 weeks of neural differentiation, performed as described [12,13], and demonstrating higher incidence of extended Tuj1$^+$ neurites elongated over 3 mm in N-CB-iPSC relative to primed CB-iPSC. (**c**) Definitive endodermal differentiation of isogenic primed vs naïve-hiPSC (*n*=6; hiPSC lines as above (**Table S1**), each individual line represented by a different color, with mean and SEM shown). Differentiations were performed as described with STEMdiff definitive endoderm kit and STEMdiff APEL medium (StemCell Technologies) [12]**.** Shown are % differentiated cells expressing FOXA2$^+$ endodermal progenitor cells. (**d,e**) Hemato-vascular (mesodermal) differentiations of non-diabetic isogenic primed vs naïve hiPSC lines were performed as described [7,12] at day 10 of embryoid body differentiation cultures for (**d**) the hESC line RUES02, and (**e**) 5 independent hiPSC lines, same as above –**Table S1**). Shown are % cells in differentiation cultures expressing surface markers for endothelial-vascular progenitors (*e.g.,* CD31$^+$, CD34$^+$, CD144$^+$, CD31$^+$CD146$^+$, KDR$^+$), pericytes (*e.g.*, CD140b$^+$, CD90$^+$NG2$^+$), angioblasts (*e.g.*, CD105$^+$, CD143$^+$), and also non-vascular lineage markers (*e.g.*, CD15$^+$). * = $p < 0.05$; ** = $p < 0.01$ (two-tailed unpaired t tests).

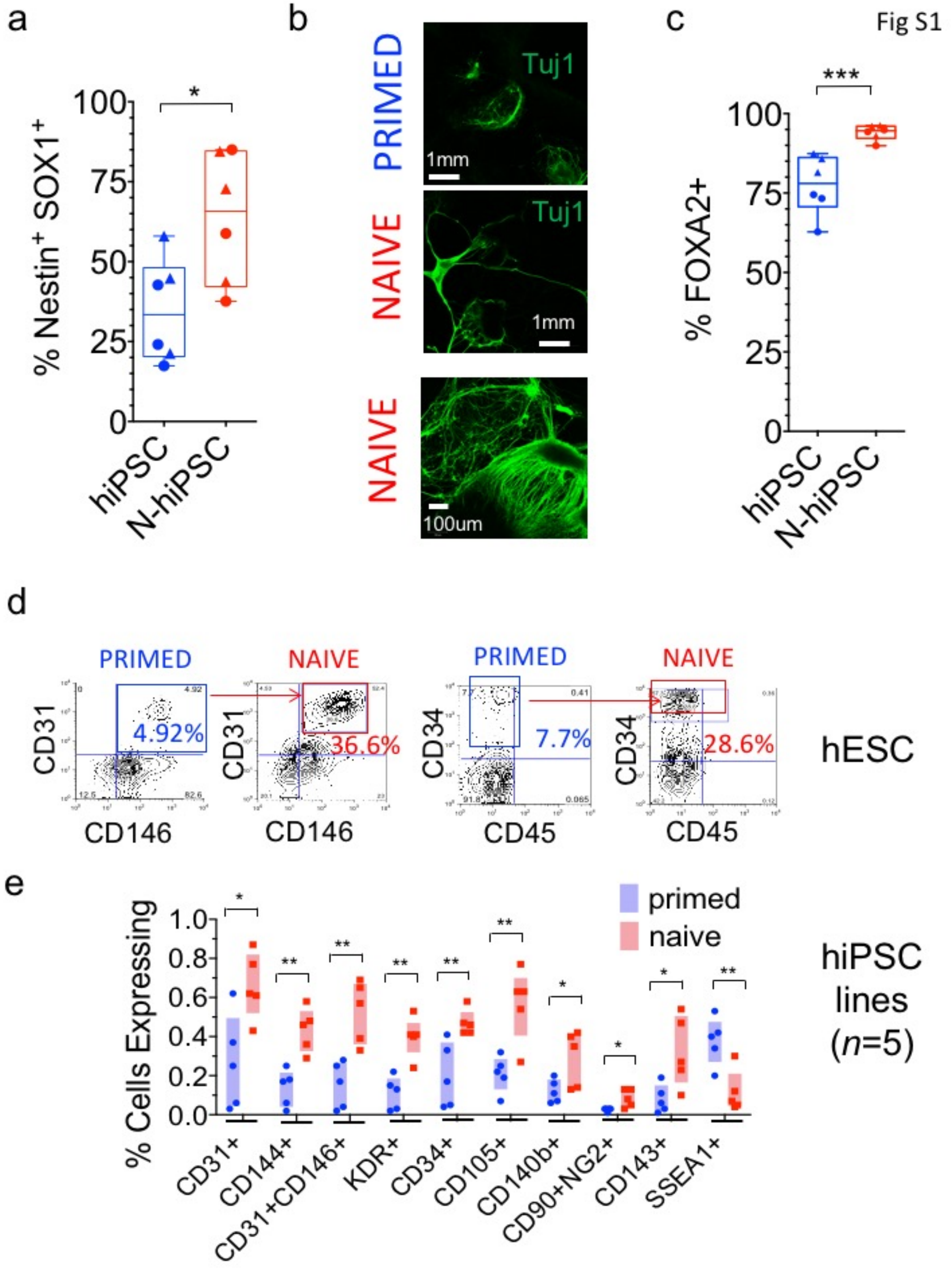

Fig S1
a
b
c
% Nestin+ SOX1+
100
75
50
25
0
*
hiPSC
N-hiPSC
PRIMED
NAIVE
NAIVE
Tuj1
Tuj1
1mm
1mm
100um
% FOXA2+
***
d
PRIMED
NAIVE
PRIMED
NAIVE
CD31
CD146
4.92%
36.6%
CD34
CD45
7.7%
28.6%
hESC
e
% Cells Expressing
1.0
0.8
0.6
0.4
0.2
0.0
primed
naive
hiPSC lines (n=5)
CD31+
CD144+
CD31+CD146+
KDR+
CD34+
CD105+
CD140b+
CD90+NG2+
CD143+
SSEA1+

**Supplementary Figure 2**. **Non-integrated episomal reprogramming of type I diabetic skin fibroblasts into conventional DhiPSC lines, and subsequent naïve reversion into N-DhiPSC with the LIF-3i culture system**. (**a**) Scheme of timeline of reprogramming of diabetic skin fibroblasts for the generation of conventional, primed DhiPSC. FGM: fibroblast growth medium, ES: ES medium, AA: ascorbic acid, CHIR99021: GSK-β inhibitor, E8: essential 8 medium. (**b**[a]) typical morphology of conventional primed DhiPSC cultured in E8 medium on vitronectin-coated plates. (**b**[b]) teratoma formation in NOG mice from primed DhiPSC line (E1CA1) generated well-developed three germ layer organoid structures. (**c**) Flow cytometry analysis of conventional DhiPSC lines demonstrated >95% $SSEA4^{+}Tra1\text{-}81^{+}$ expressions. (**d**) Scheme of timeline of naïve reversion (LIF-5i/LIF-3i) of primed, conventional DhiPSC into N-DhiPSC, performed as described [12]. (**e**) typical dome-shaped colonies (left, [a,c]) in LIF-3i naïve cultures of N-DhiPSC lines (E1C1, E1CA2 – see **Table S1**) and their normal G-banded karyotypes (right, [b,d]) following 3-10 passages in naïve (LIF-3i) conditions. (**f**) Representative teratoma H&E sections from two N-DhiPSC lines (E1CA2 and E1C1). Shown are ectodermal neural rosette (Ect), endodermal epithelial gut (End), and mesodermal cartilage (Meso) structures. Scale bar = 100 um.

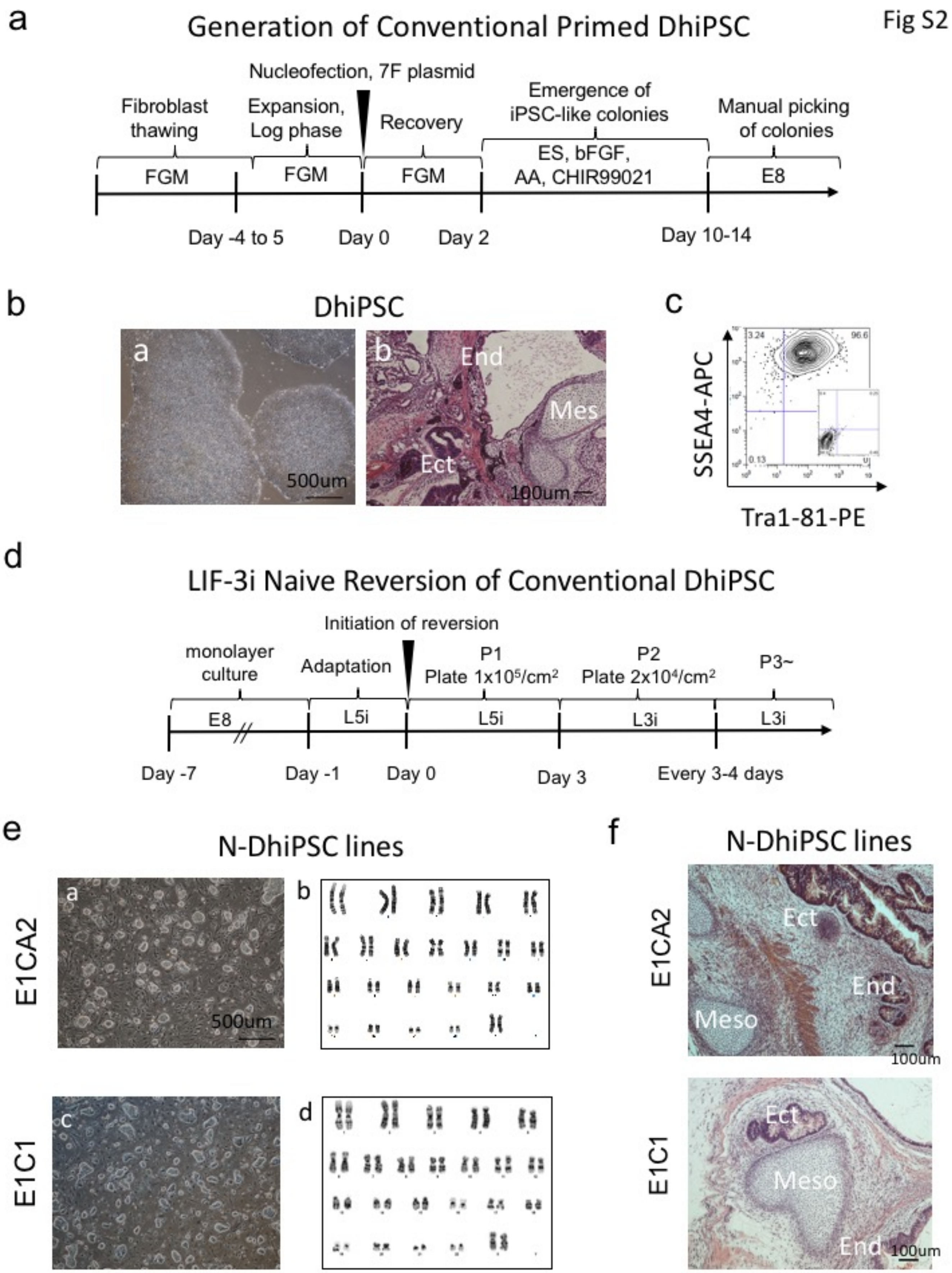
a
Generation of Conventional Primed DhiPSC
Fig S2
Nucleofection, 7F plasmid
Fibroblast thawing
Expansion, Log phase
Recovery
Emergence of iPSC-like colonies
Manual picking of colonies
FGM
FGM
FGM
ES, bFGF, AA, CHIR99021
E8
Day -4 to 5
Day 0
Day 2
Day 10-14
b
DhiPSC
a
b
End
Mes
Ect
500um
100um
c
SSEA4-APC
Tra1-81-PE
3.24
96.6
0.13
d
LIF-3i Naive Reversion of Conventional DhiPSC
Initiation of reversion
monolayer culture
Adaptation
P1 Plate 1x10⁵/cm²
P2 Plate 2x10⁴/cm²
P3~
E8
L5i
L5i
L3i
L3i
Day -7
Day -1
Day 0
Day 3
Every 3-4 days
e
N-DhiPSC lines
E1CA2
E1C1
a
b
c
d
500um
f
N-DhiPSC lines
E1CA2
E1C1
Ect
End
Meso
Ect
Meso
End
100um
100um

**Supplementary Figure 3**. **APEL vascular differentiation of primed and naïve DhiPSC**. (**a**) Both primed (E8) and naïve (LIF-3i) DhiPSC (line E1C1) differentiated efficiently using the APEL monolayer vascular differentiation system. Shown are morphologies of APEL-differentiated DVP cells before and after CD31-sorting and re-plating in EGM2. (**b**) Representative flow cytometry expressions of CD31 and CD146 prior to and post sorting of APEL differentiation cultures with CD31 magnetic bead-tagged antibody (MACS: magnetic activated cell sorting). (**c**) Post CD31-sort (prior to EGM2 expansion) vascular lineage surface marker analysis of primed DVP (left panel) vs N-DVP (right panel) demonstrating no significant differences in vascular marker expressions post CD31-sorting from APEL cultures. (**d**) $CD31^{+}CD146^{+}$ N-VP and N-DVP were generated from normal non-diabetic naïve N-CBiPSC and naïve N-DhiPSC, respectively, and analyzed post-CD31 sorting for surface expressions of vascular markers (*e.g.*, CD31, CD146, CD144, CXCR4, CD90, and CD105); which demonstrated no significant differences.

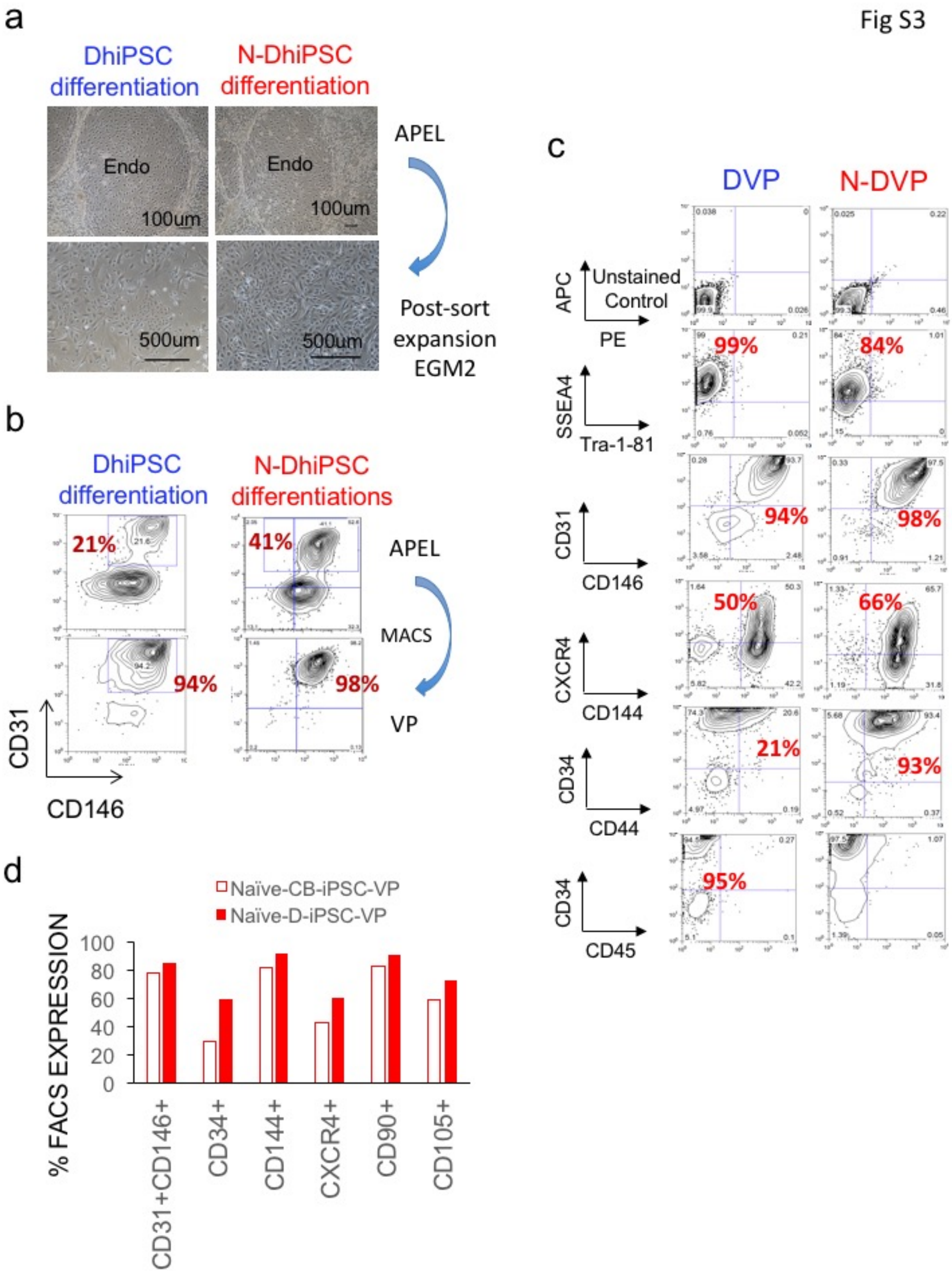

a
Fig S3
DhiPSC differentiation
N-DhiPSC differentiation
Endo
Endo
100um
100um
500um
500um
APEL
Post-sort expansion EGM2
b
DhiPSC differentiation
N-DhiPSC differentiations
21%
41%
94%
98%
APEL
MACS
VP
CD31
CD146
c
DVP
N-DVP
APC
Unstained Control
PE
99%
84%
SSEA4
Tra-1-81
94%
98%
CD31
CD146
50%
66%
CXCR4
CD144
21%
93%
CD34
CD44
95%
CD34
CD45
d
Naïve-CB-iPSC-VP
Naïve-D-iPSC-VP
% FACS EXPRESSION
100
80
60
40
20
0
CD31+CD146+
CD34+
CD144+
CXCR4+
CD90+
CD105+

**Supplementary Figure 4**. ***In vitro* vascular function of primed DVP vs. N-DVP.** (**a**) Representative staining images of β-galactosidase senescent cell assay at passage 4 post re-plating of primed VP vs. N-VP from a normal fibroblast-hiPSC (C1.2), and primed DVP vs N-DVP from a diabetic hiPSC line (E1C1). (**b**) Representative flow cytometry analysis of EdU assays at passage 1 and passage 3 post re-plating of primed DVP vs. N-DVP from a diabetic hiPSC line (E1C1) (see **Figure 4e**). (**c**) Matrigel vascular tube formation assay demonstrating that N-DVP formed longer and more mature types of tubes than primed DVP.

Fig S4

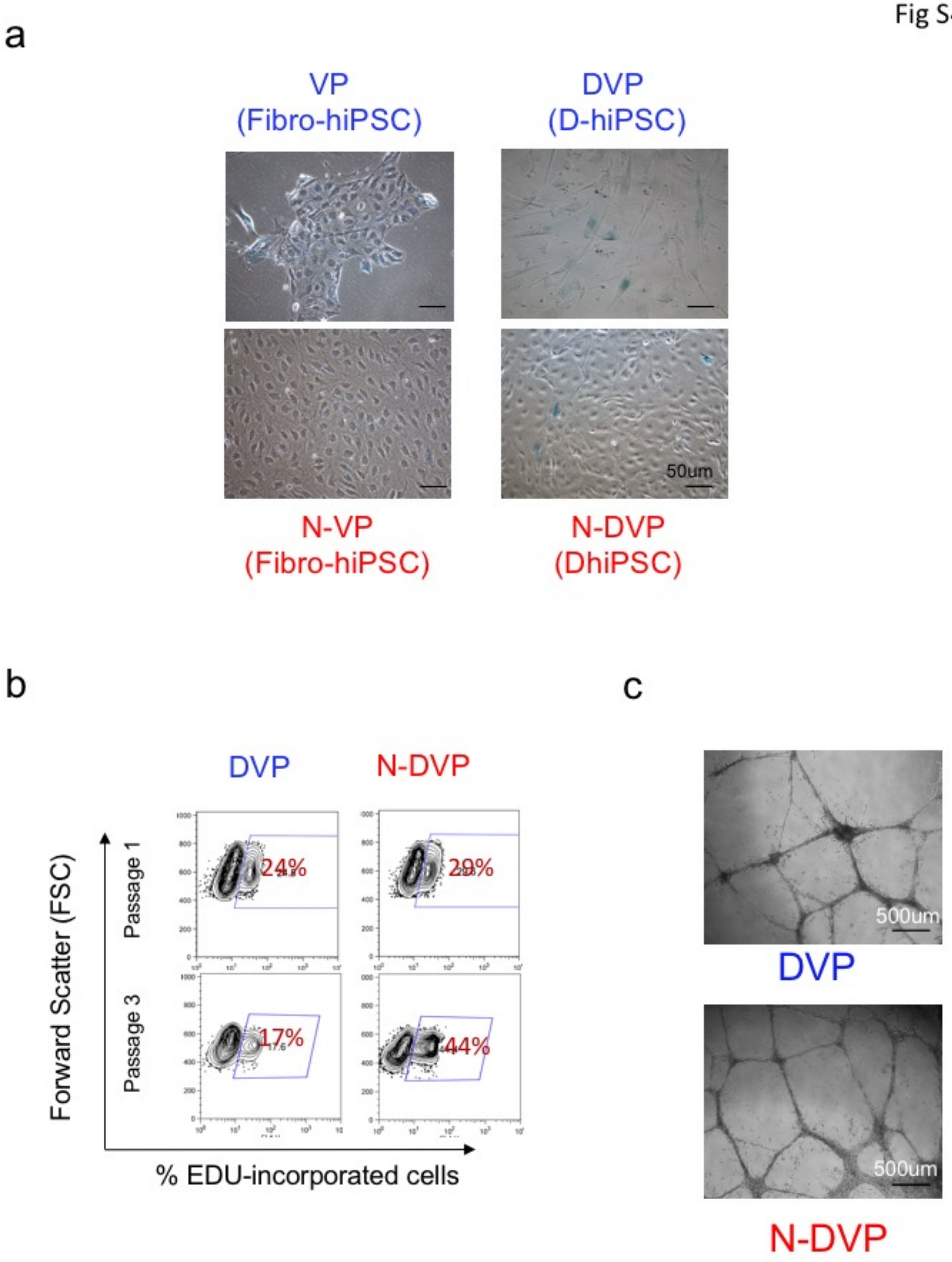
a
VP
(Fibro-hiPSC)
DVP
(D-hiPSC)
50um
N-VP
(Fibro-hiPSC)
N-DVP
(DhiPSC)
b
DVP
N-DVP
Forward Scatter (FSC)
Passage 1
Passage 3
24%
29%
17%
44%
% EDU-incorporated cells
c
500um
DVP
500um
N-DVP

**Supplementary Figure 5**. **Western blot analysis of DNA damage response (DDR) proteins following treatment of primed VP and N-VP with NCS.** Purified and re-plated DVP vs N-DVP differentiated from isogenic primed DhiPSC *vs*. N-DhiPSC (line E1C1) or VP vs N-VP differentiated from isogenic primed vs naïve normal fibroblast-hiPSC (C2); with (+) and without (-) NCS treatment prior cell lysate collection. P-DNA-PK: phosphorylated DNA-PK; P-H2AX: phosphorylated H2AX; NCS: neocarzinostatin.

Fig S5

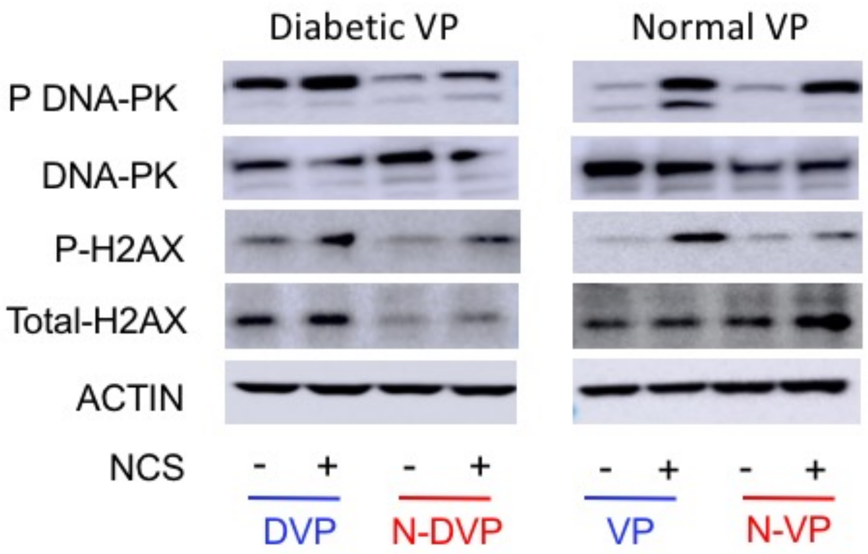

Diabetic VP
Normal VP
P DNA-PK
DNA-PK
P-H2AX
Total-H2AX
ACTIN
NCS
- + - +
- + - +
DVP
N-DVP
VP
N-VP

**Supplementary Figure 6**. **Human VP cell therapy of murine ischemic retinopathy following ischemia-reperfusion (I/R) injury**. Representative high magnification images from whole mount retinae demonstrating more abundant survival of $HNA^{+}$ N-DVP cells in the superficial vascular layers of the retina relative to primed DVP at (**a**) 7 days and (**b**) three weeks following parallel intra-vitreal injections of 50,000 DVP or N-DVP cells per eye. (**b**) Retinal vascular regions at 3 weeks demonstrated that $HNA^{+}$ human DVP cells were located either abundantly in suspension within vitreous or had migrated and engrafted into vascular abluminal regions of murine ischemic vessels (arrow heads). Scale bars = 50 μm. Whole mount retinae were also stained with anti-murine CD31 (mCD31) or anti-murine collagen IV (mColV) antibodies for z-stack and maximum-intensity projection imaging. (**c,d**) Human vascular engraftment of murine retinae, at 2 weeks post DVP injection. (**c**) Whole mount retinal staining of mouse CD31 (mCD31) and co-staining with murine collagen type-IV (mCol-IV) at 2 weeks following I/R injury and naïve (N-DVP) injections; (**d**) co-staining with human CD34 (hCD34)-specific and murine collagen type-IV (mCol-IV) antibodies of similar whole mount retinal samples demonstrated that a subset of these mCD31+mCol+ murine vessels had also formed CD34+ human-murine vascular chimerism.

Fig S6

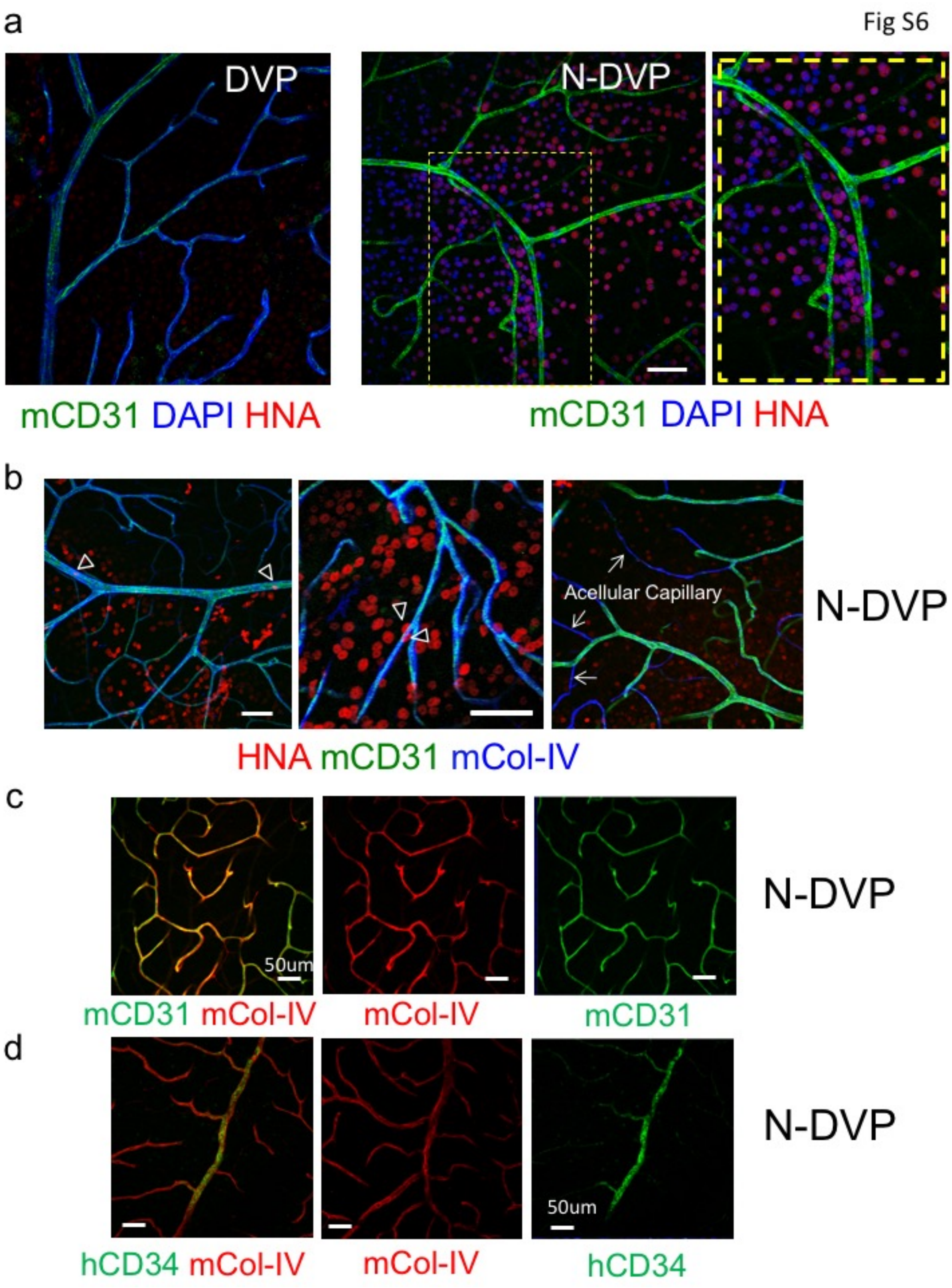
a
DVP
N-DVP
mCD31 DAPI HNA
mCD31 DAPI HNA
b
Acellular Capillary
N-DVP
HNA mCD31 mCol-IV
c
50um
N-DVP
mCD31 mCol-IV
mCol-IV
mCD31
d
50um
N-DVP
hCD34 mCol-IV
mCol-IV
hCD34

**Supplementary Figure 7**. **Method of quantitating human DVP cell engraftment in the vasculature of the neural layers of ischemia-injured mouse retinae**. (**a**) Tile-scanned image of cryo-sectioned retina. Multiple images were taken in 20x magnification from each cryo-sections as well as spaced (serial sectioned) retina sections for quantification studies. Hatched areas demonstrate representative example of neural retinal regions evaluated for human vascularization of murine blood vessels. (**b**) Method of regional separation of neural retinal layers employed for quantification of human $CD34^+$ cells (hCD34) for data presented in **Figure 7b** and **7d** via automated counts using FiJi distribution of ImageJ software.

Fig S7

a

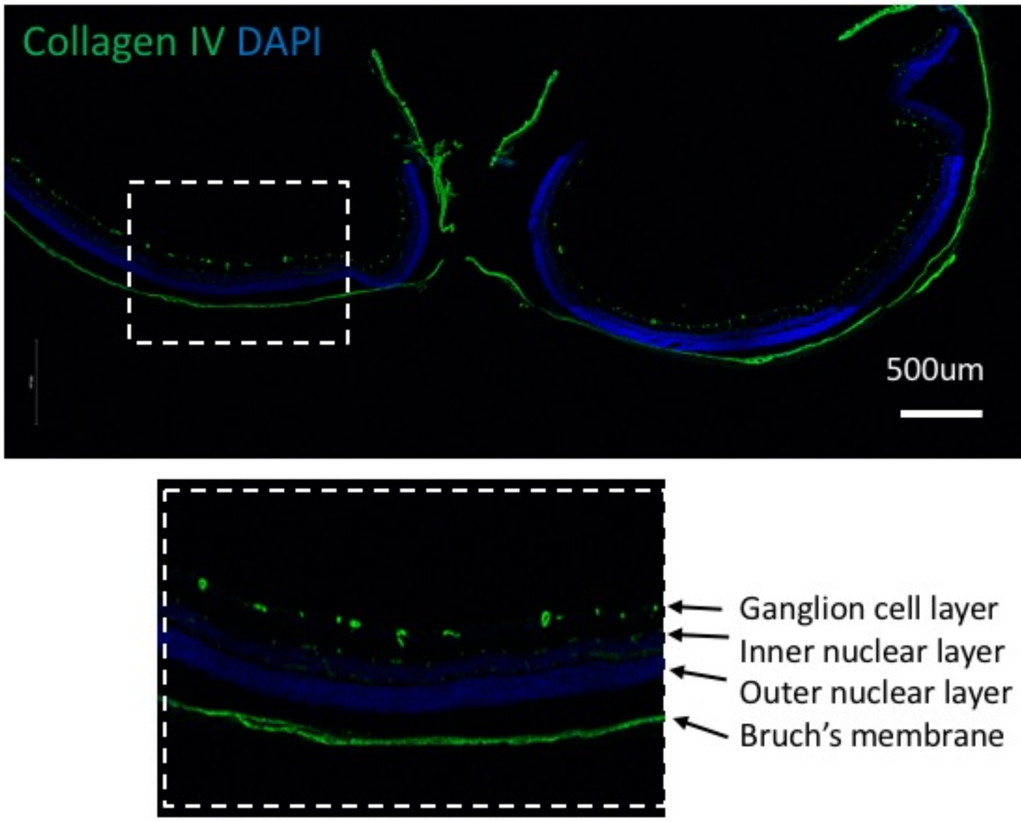


b

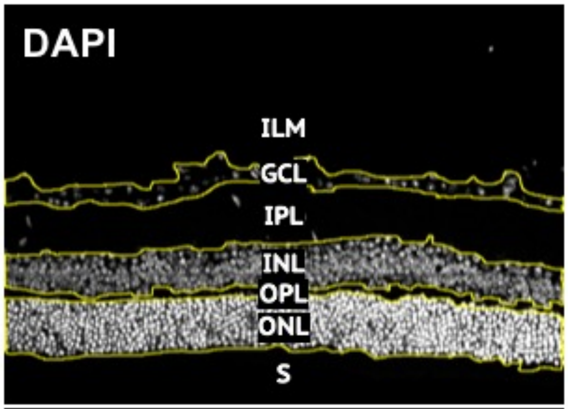


| | |
|---|---|
| inner limiting membrane | ILM |
| ganglion cell layer | GCL |
| inner plexiform layer | IPL |
| inner nuclear layer | INL |
| outer plexiform layer | OPL |
| outer nuclear layer | ONL |
| segments | S |

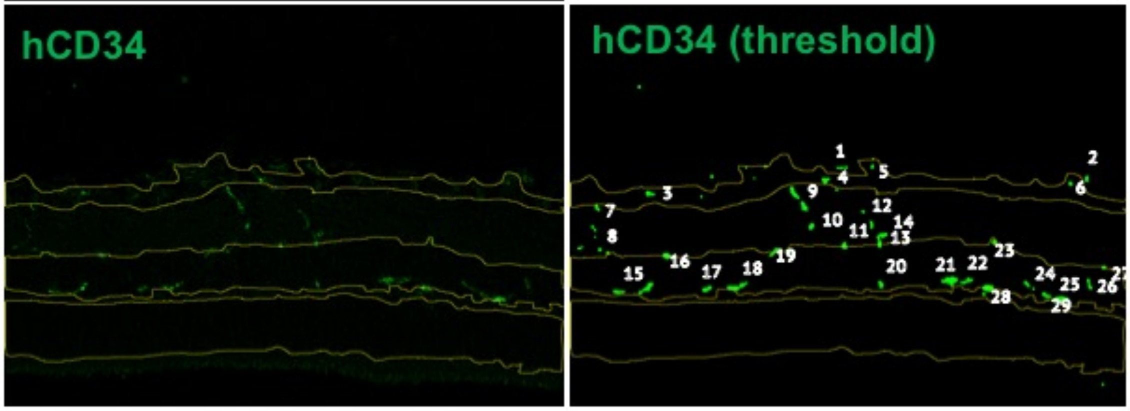

**Supplementary Figure 8**. **Lineage-primed gene expression and epigenetic configurations of PRC2-regulated bivalent promoters in primed vs naïve DhiPSC**. (**a**) Crossplot of mean genome-wide gene expression of PRC2 module gene targets (**Table S3**) *vs.* CpG methylation of non-diabetic N-hiPSC *vs.* their isogenic primed isogenic hiPSC counterparts (*n*=6 normal hiPSC lines; **Table S1**). Plotted are the differentially methylated region (DMR) CpG methylation beta values of PRC2 module promoter regions in LIF-3-reverted hiPSC minus their isogenic primed hPSC counterparts (y-axis, p ≤0.05) *vs*. their corresponding differential gene expressions for the same genes (red, x-axis, $log_2$ fold changes (FC); p ≤ 0.05, FC +/- ≥1.5.). **(b)** Curated GSEA pathways for gene targets of the PRC2 module over-represented (FDR<0.01; p<0.001) in LIF-3i-reverted fibroblast-hiPSC *vs*. their isogenic primed counterparts (*n*=4; normal hiPSC lines –**Table S1**). (**c**) Comparison of differentially expressed (p ≤ 0.05, fold change (FC) +/- ≥1.5) lineage-primed transcriptional targets of the PRC2 module (**Table S3**) in naïve vs primed hiPSC (*n*=6 independent lines). FC: normalized ratios of naïve/primed expression microarray signal intensities show broadly decreased expressions of PRC2 targets in N-hiPSC lines. Ratios are of LIF-3i-reverted *vs.* primed hPSC samples. (**d**) CHiP-qPCR for H3K27me3 and H3K4me3 histone marks at key multi-lineage developmental promoters in primed vs N-DhiPSC line E1C1. Data is presented as percent input materials between naïve and primed D-hiPSC line E1C1. Standard error of the mean (SEM) bar represents replicates. ChiP-PCR primer sequences and sources are detailed in **Table S3**.

.

Fig S8

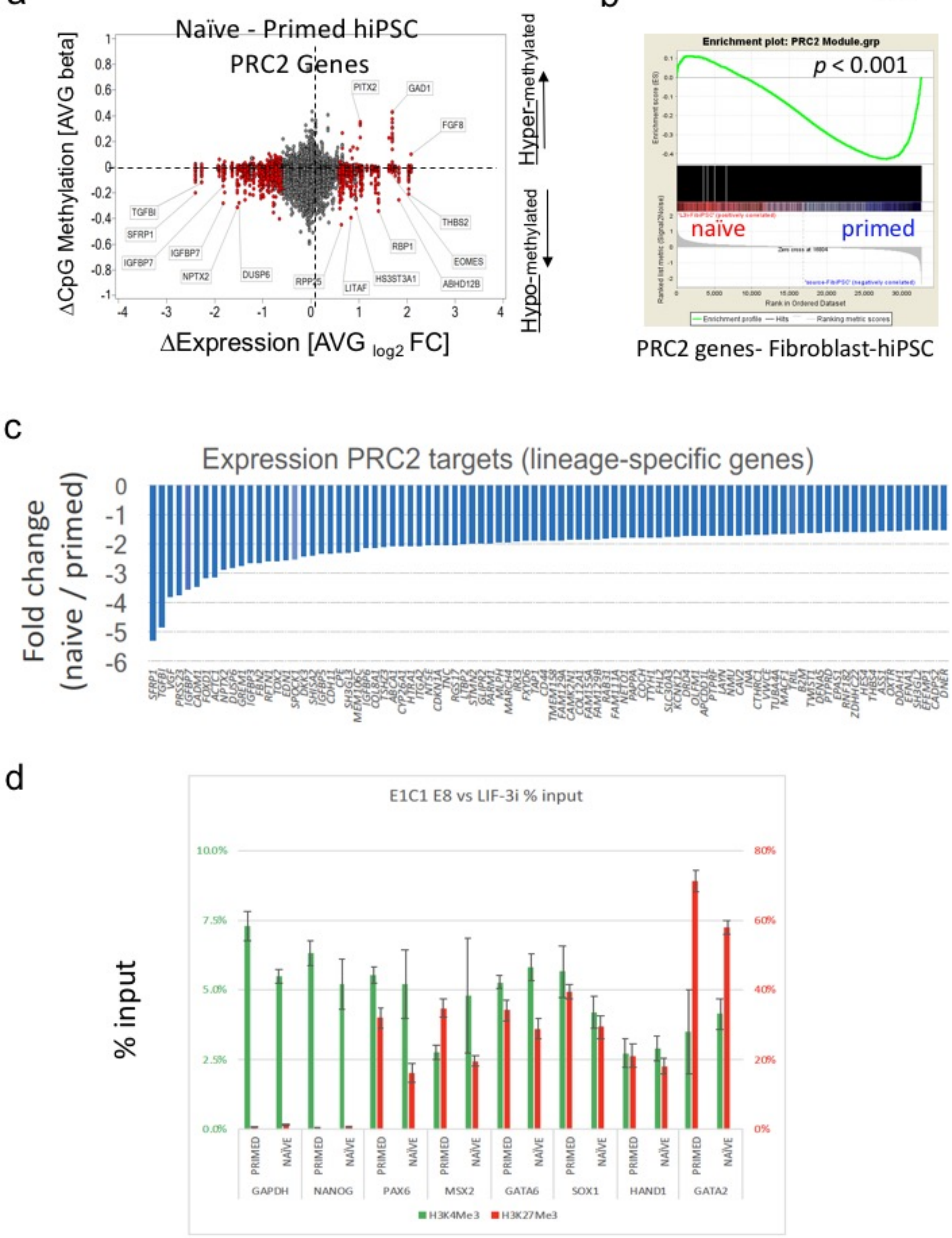
a
Naïve - Primed hiPSC
PRC2 Genes
ΔCpG Methylation [AVG beta]
ΔExpression [AVG log2 FC]
Hyper-methylated
Hypo-methylated
PITX2
GAD1
FGF8
TGFBI
SFRP1
IGFBP7
IGFBP7
NPTX2
DUSP6
RPP25
LITAF
HS3ST3A1
RBP1
THBS2
EOMES
ABHD12B
b
Enrichment plot: PRC2 Module.grp
p < 0.001
naïve
primed
Rank in Ordered Dataset
Enrichment profile
Hits
Ranking metric scores
PRC2 genes- Fibroblast-hiPSC
c
Expression PRC2 targets (lineage-specific genes)
Fold change (naive / primed)
d
E1C1 E8 vs LIF-3i % input
% input
PRIMED
NAÏVE
GAPDH
NANOG
PAX6
MSX2
GATA6
SOX1
HAND1
GATA2
H3K4Me3
H3K27Me3
